# Recolonisation strategies of early animals in the Avalon (Ediacaran 574 – 560 Ma)

**DOI:** 10.1101/2025.01.30.635654

**Authors:** Nile P. Stephenson, Katie M. Delahooke, Princess A. Buma-at, Benjamin W. T. Rideout, Nicole Barnes, Charlotte G. Kenchington, Andrea Manica, Emily G. Mitchell

**Affiliations:** Department of Zoology, University of Cambridge, UK; University Museum of Zoology, University of Cambridge, UK; Department of Earth Sciences, University of Cambridge, UK; Independent

**Keywords:** Ediacaran, secondary succession, spatial ecology, Avalon, *Fractofusus*

## Abstract

The first geographically widespread metazoans are found in the Avalon assemblage (Ediacaran; 574 – 560 Ma). These early animals were regularly disturbed by sedimentation events such as ash flows and turbidites, leading to an apparent “resetting” of communities. However, it is not clear how biological legacies – remains or survivors of disturbance events – influenced community ecology in the Avalon. Here, we use spatial point process analysis on 19 Avalon palaeocommunities to test whether two forms of biological legacy (fragmentary remains of *Fractofusus* and surviving frondomorphs) impacted the recolonisation dynamics of Avalon palaeocommunities. We found that densities of *Fractofusus* were increased around the *Fractofusus* fragments, suggesting that they helped to recolonise the post-disturbance substrate, potentially contributing to the *Fractofusus* dominance found in 8 of the 19 palaeocommunities. However, we found no such effects for survivor fronds. Our results suggest that the evolution of height was for long-distance dispersal rather than local recolonisation. In modern deep-sea environments, there is a trade-off between local and long-distance dispersal, and our work demonstrates that this differentiation of reproductive strategies had already developed in the early animals of the Avalon.

## Introduction

The first geographically widespread metazoans dominate the Avalon assemblage (574 – 560 million years ago) during the terminal Ediacaran (Dunn et al., 2021; Liu et al., 2015; Mitchell & Pates, 2025; Runnegar, 2022). Avalon palaeocommunities were dominated by enigmatic rangeomorphs and arboreomorphs (Brasier et al., 2012; Narbonne, 2004), alongside rare, more familiar crown-group cnidarians (Dunn et al., 2022) and putative sponges (Aragonés Suarez & Leys, 2022; Sperling et al., 2011). The Avalon palaeocommunities are preserved *in situ* and near-census (O’Brien & King, 2005) in Newfoundland, Canada, and Charnwood Forest, Leicestershire, UK (Benus, 1988; Noble et al., 2015; Wood et al., 2003), in deep water, slope environments adjacent to a volcanically-active island arc system (Wood et al., 2003). Disturbance from sedimentation events (ash influx, turbidites) was likely to be a substantial and potentially regular cause of mortality for these early animal communities (Clapham et al., 2003; Wilby et al., 2015). Environmental disturbance has therefore been suggested to have substantially shaped the ecology and evolution of Avalon communities (Clapham et al., 2003; Delahooke et al., 2024; Kenchington et al., 2018; Wilby et al., 2015).

Periodic pulse disturbances by intensive sedimentary events smothered entire Avalonian communities, and so were typically lethal (O’Connell et al., 2024; Wilby et al., 2015), leaving behind barren substrate. For new communities to emerge, re-colonisation processes are required. These can be either a new primary succession, sourced via long-distance dispersal from nearby surviving communities (Boddy et al., 2022; Clapham et al., 2003; Eden et al., 2022; Mitchell et al., 2015), or a secondary succession, from local recolonisation whereby either biological remains or surviving organisms aid local recolonisation – a “biological legacy” (Wilby et al., 2015). In modern sessile systems, surviving organisms often have a substantial impact on the post-disturbance community composition (Bače et al., 2015), ecosystem functions (Nyström & Folke, 2001), and ecological dynamics (Wild et al., 2014). Usually, survivors represent the most common taxa in a given area (Wild et al., 2014), generating legacy effects which can determine the spatial ecology of the recolonising community, where the secondary community is characterised either by clustering around suitable habitat near biological remains (e.g., logs and tree stumps), or by reproductive clusters around survivors (e.g., coral fragments) (Bače et al., 2015; Kim et al., 2022; Maldonado & Uriz, 1999; Okubo et al., 2007; Wild et al., 2014). Such legacies can sometimes effect succession trajectories and lead to alternative stable states (Jõgiste et al., 2017; Pérez-Hernández & Gavilán, 2021). However, such ecological benefits of biological legacies for Avalon palaeocommunities could be compounded by the relative slowness of re-colonising processes because of their deep-sea habitat, similar to the slowness of modern deep-sea recolonising processes (Grassle, 1977). Still, the temporal record of many Avalonian taxa stretches over millions of years, demonstrating that the taxa could survive regular disturbances.

Within Avalon palaeocommunities, there are two possible examples of specimens that could provide evidence of biological legacies. First, fragmented fossils of *Fractofusus andersoni* (Fig. 1a) alongside complete, but much smaller *Fractofusus andersoni* fossils on the Brasier surface in Mistaken Point Ecological Reserve, Newfoundland, Canada could represent vegetative, reproductive fragments - perhaps generated by a disturbance event. Secondly, surviving individuals could be identified as an outlier(s) within their population size distribution in the post-disturbance community (Wilby et al., 2015) (Fig. 1b-d) because any individuals surviving the disturbance event would be older and therefore taller than individuals of the post-disturbance community. Such outlier individuals have been observed for *Primocandleabrum, Charnia, Hyllaecullulus,* and *Charniodiscus* in North Quarry Bed B (henceforth, Bed B) in Charnwood Forest (Wilby et al., 2015). The advantages of local reproduction following the survival of disturbance events could therefore provide an additional driver for the evolution of longer-distance dispersal (Mitchell & Kenchington, 2018).

**Figure 1:**
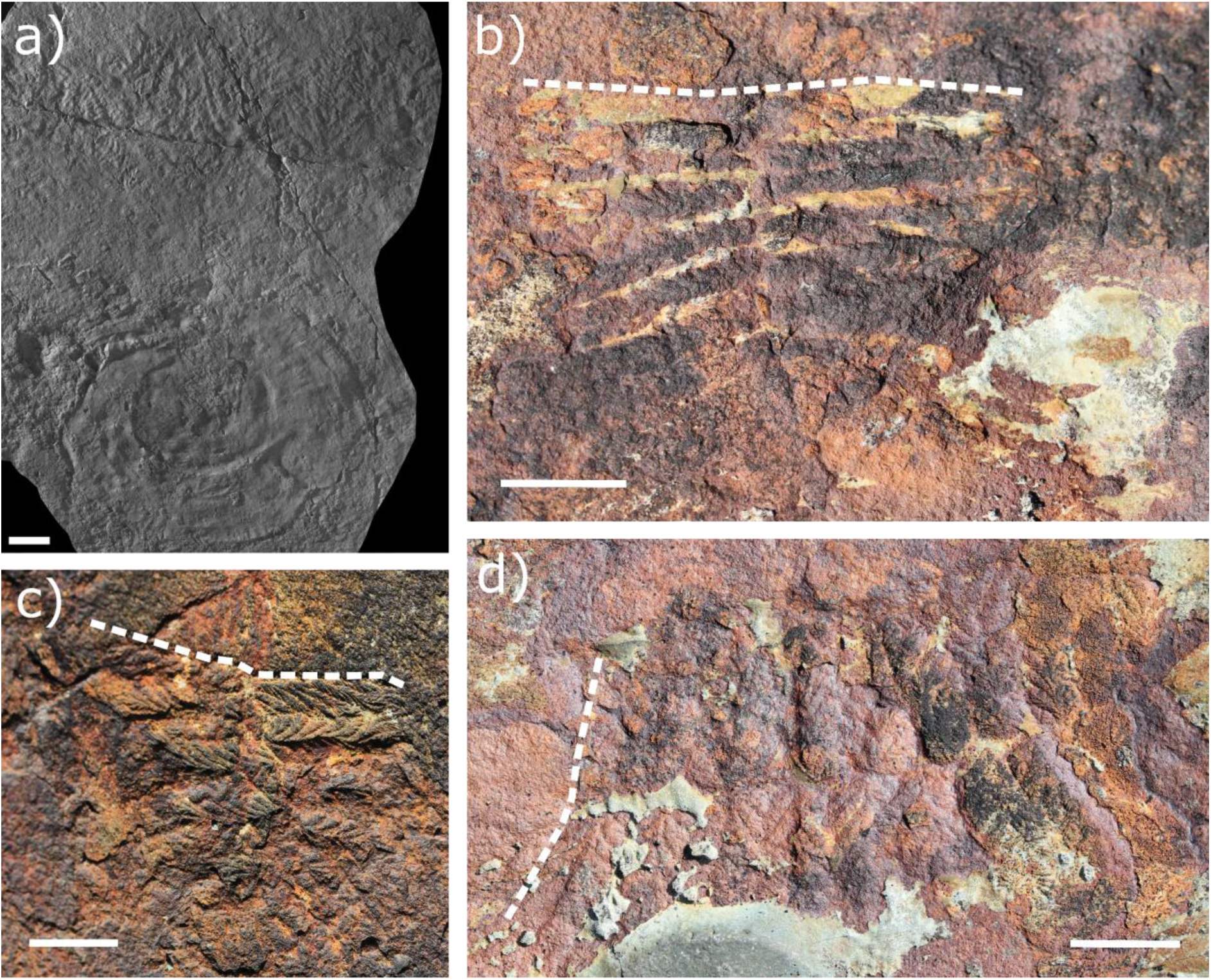
a) Outlier *Primocandelabrum* sp. on Bed B. Scale bar = 2 cm. b) – d) fragment *Fractofusus andersoni* specimens from the Brasier surface. Dashed lines denote broken edges of specimens. Scale bars = 1 cm.

Spatial point process analysis enables the inference of ecological processes from the underlying spatial patterns of points (e.g., fossils), because for sessile organisms, such as Avalonian taxa, there is only a limited set of possible processes behind observed spatial patterns. These limited set of processes mean that by comparing the observed patterns with those produced by well-established models, we can determine the most likely underlying processes for any patterns found across plant, coral, fungi, and sessile fossil communities (McFadden et al., 2019; Mitchell & Harris, 2020; Velázquez et al., 2016). Therefore, these techniques can be used to investigate the potential influences of re-colonisation processes in Avalon palaeocommunities. We hypothesise: i) that *Fractofusus andersoni* fragments on the Brasier surface have a substantial positive effect on *Fractofusus* population structure (densities in proximity to fragments redolent of a reproductive mechanism); and ii) that the presence of outlier individuals has a substantial positive effect on community structure (relative abundances) and population spatial structure (higher densities in proximity to the outlier specimens) which we systematically test using SPPA.

### Geological setting

Avalonian Ediacaran macrofossils occur in the Conception and St. John’s groups in Newfoundland, Canada, and the Charnian Supergroup in Leicestershire, UK, specifically within thick siliciclastic-volcaniclastic units comprised of fine-grained turbiditic facies. These successions represent deep-marine slope environmental settings (Myrow, 1995; O’Brien et al., 1983), and substantial volcaniclastic sediments in these units indicates ash input from nearby volcanic arcs (Wood et al., 2003). The palaeocommunities in this study come from the Mistaken Point Ecological Reserve (MPER) (Benus, 1988) (Fig. 2), Bay Roberts and Spaniard’s Bay (Ichaso et al., 2007), and the Discovery Geopark (DG) in the Bonavista Peninsula (Hofmann et al., 2008). The MPER and DG successions were likely deposited in different deep-water basins within the Avalonian Terrane (O’Brien & King, 2005).

**Figure 2:**
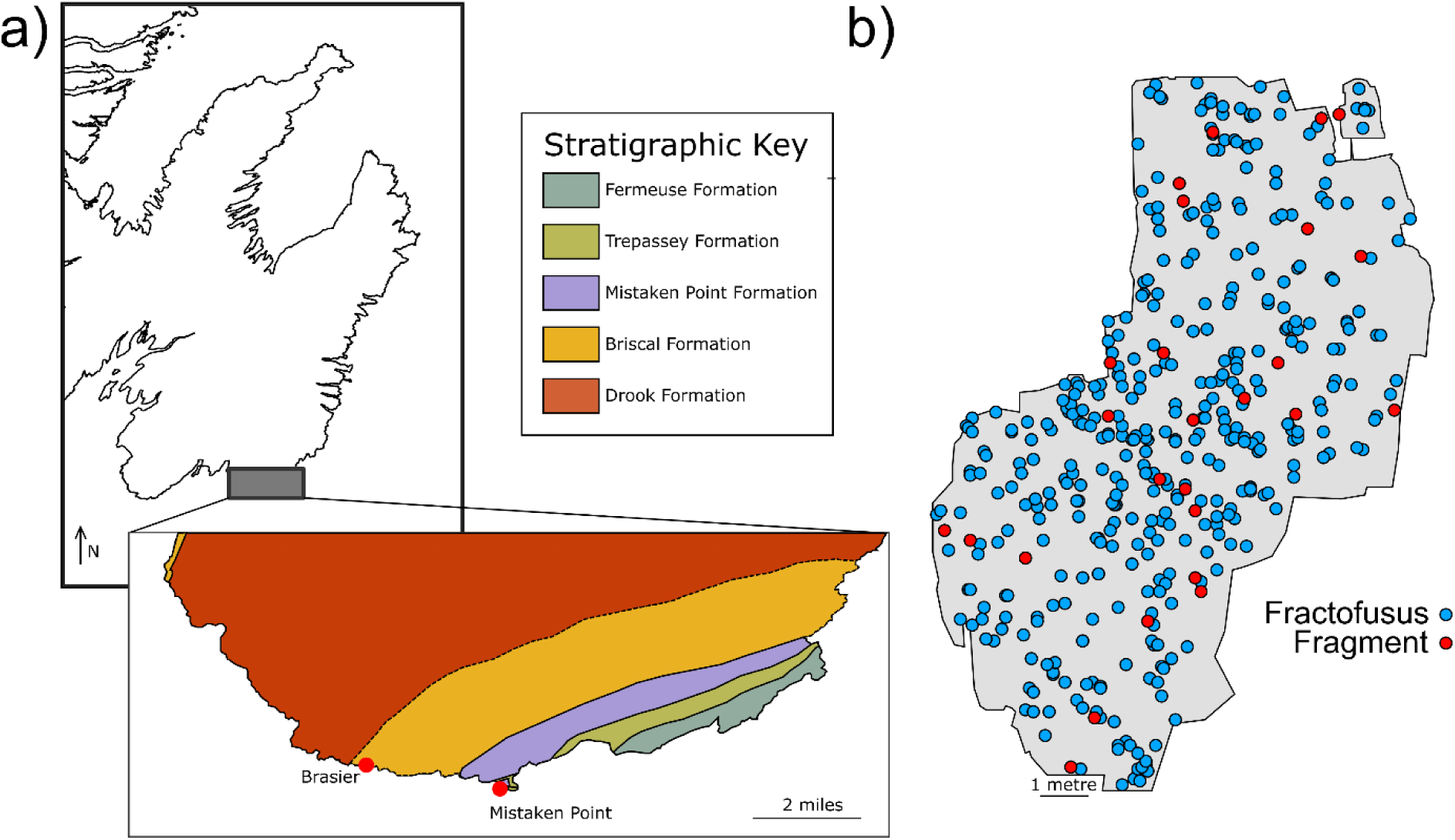
a) Stratigraphic map of Mistaken Point Ecological reserve (adapted from (Matthews et al., 2021)), with Brasier surface labelled. b) Brasier surface with complete (blue) and fragmented (red) *Fractofusus andersoni* specimens.

The three key surfaces within this study are i) The Brasier Surface, ii) the E Surface, and iii) Bed B. The Brasier Surface (567.8 Ma, Fig. 2), which occurs within the lower Briscal Formation, consists of normally graded medium- to coarse-grained sandstones with a greenish hue due to their high tuffaceous content (Matthews et al., 2021). The E surface (565 Ma), within the Mistaken Point Formation, is characterised by medium-bedded siltstones deposited by strong turbidity flows, capped by fine-grained hemipelagic sediments containing coarse-grained crystal tuffs (Narbonne, 2005). Bed B, Charnwood Forest is within the Bradgate Formation, which consists of fine-grained sedimentary-volcaniclastic rocks deposited in deep-marine slope settings, well below the storm wave base, within a back-arc basin (Le Bas, 1984; Pharaoh et al., 1987). The base of the Bradgate Formation is dated to be 561.9 ± 0.9 Ma, so provides a lower bound for the age of Bed B (Noble et al., 2015).

## Materials and Methods

### Material

In this study, we assessed 19 bedding plane assemblages: “Yale” outcrops of the D and E surfaces at Mistaken Point; and the Bristy Cove surface from the Mistaken Point Ecological Reserve; the St. Shott’s surface at Western Head, Newfoundland, Canada from (Mitchell et al., 2019); the Brasier surface; the Clapham’s Pigeon Cove (CPC) surface; the Shingle Head surface; the Lower Mistaken Point surface; the “Queens” outcrop of the G Surface at Mistaken Point; and the Pizzeria surface from the Mistaken Point Ecological Reserve, Newfoundland, Canada using data from (Stephenson et al., 2024); the Bishop’s Cove surface and the Green Point surface from the Bay Roberts area in Newfoundland, Canada from (Stephenson et al., 2024); the H14 surface from (Mitchell et al., 2019), the H5 surface from (Delahooke et al., 2024), and the H26 surface, the Capelin Gulch site, the Goldmine surface, and the H38 surface from (Stephenson et al., 2024) all in the Discovery Geopark in Newfoundland, Canada; and Bed B from Charnwood Forest from (Mitchell et al., 2019).

The surfaces were mapped using a combination of methods *cf* (Mitchell et al., 2019). All surfaces were LiDAR scanned using (Faro Focus^m^, mean resolution up to 0.5 mm) (apart from the H38 surface, which was not scanned) and mapped using photogrammetry in Agisoft Metashape v1.7.5. Additional laser-line probe scans were used for the Pizzeria, Brasier, G, D, and E surfaces to a 0.050 mm resolution (Mitchell et al., 2019). In order to create the specimen maps, three-dimensional models created from photogrammetry were used to make orthomosaics (two-dimensional photomaps) from which two-dimensional vector maps containing information on the spatial position, disc length and width, stem and frond length and width, and species-level identification were made *cf* (Stephenson et al., 2024) (Extended Data Table 3) in Inkscape v1.3. *Fractofusus* fragments were specifically noted within the Brasier surface dataset. A custom script (https://github.com/nis38/NPS_dex in R v 4.2.2 (R Core Team, 2023) adapted from (Delahooke et al., 2024)) was used to extract data from vector maps. All surface datasets (surface outlines, specimen sizes, specimen spatial positions) were retrodeformed by quantifying tectonic deformation through comparisons of the length and widths of holdfasts >10 mm, with the assumption of original circularity (Wood et al., 2003), apart from on the CPC surface, which did not have sufficient holdfasts (Clapham et al., 2003). The total dataset comprised 18,060 specimens from 43 morphotaxa (taxa listed in Table S2 plus effaced fronds, discs, and ivesheadiomorphs).

### Spatial analysis of fragments

In this study, we investigated the population ecology of *Fractofusus* (whole and fragmented specimens) on the Brasier surface (Fig. 1b-d) by means of univariate and bivariate spatial point process analysis (SPPA) (Mitchell et al., 2019; Wiegand & Moloney, 2014). In order to compare the differences in size between fragmented and complete *Fractofusus* specimens, we used a Mann-Whitney U test (Mann & Whitney, 1947).

We used the pair correlation function (PCF) summary statistic to quantify how the density of points (here, fossils) at a given distance *r* from a given point change as a function of distance from that point (Illian et al., 2008; Wiegand & Moloney, 2014). The simplest spatial pattern, complete spatial randomness (CSR), can be modelled as a homogenous Poisson process (Illian et al., 2008). CSR reflects no biotic or abiotic interactions at the spatial scales considered (Illian et al., 2008; Velázquez et al., 2016). Where spatial patterns are not CSR, they can be aggregated (PCF > 1) or segregated (PCF < 1), and these patterns can be detected at a range of spatial scales and combinations of aggregations and segregations (Illian et al., 2008). Aggregation within a population can be caused by habitat associations, whereby organisms cluster within an area of preferable resources (*cf* (Shen et al., 2009)), by dispersal-limited reproduction (*cf* (McFadden et al., 2019; Mitchell et al., 2015)), or by both processes contemporaneously (Wiegand & Moloney, 2014). Habitat-associated aggregation processes can be modelled by heterogeneous Poisson (HP) processes (Wiegand & Moloney, 2014), whereas reproductive aggregation can be modelled by Thomas cluster (TC) processes (Mitchell et al., 2015; Thomas, 1949; Wiegand & Moloney, 2014). Reproductive- and habitat-associated processes combined would thus be best modelled by Thomas clustering processes over a heterogeneous background – inhomogeneous Thomas clusters (ITC) (Illian et al., 2008).

In order to understand if complete *Fractofusus* specimens were clustered around fragments, a density map of the fragments using an Epanechnikov kernel (Wiegand & Moloney, 2014) and a distance of 1.7 m was produced. A kernel of 1.7m was used because this distance provided a balance between the locations of the fragments and the variation generated by them. This density map was used as a covariate in HP and ITC models (Illian et al., 2008) in Programita (v February 2014) (Wiegand, 2014). CSR and HP models were fit using maximum likelihood methods and TC and ITC models were fit using minimum contrast methods (Baddeley, 2015; Diggle & Gratton, 1984). To test whether a given PCF was best fit to any given theoretical model PCF, 9,999 Monte Carlo simulations were run for the theoretical process, and the simulation envelopes chosen to be between the 5% highest and lowest values (Baddeley, 2015).

In order to investigate the differences in the spatial patterns between the whole versus fragment *Fractofusus* specimens, we used random labelling analysis (RLA) (Getzin et al., 2006, 2008). RLAs are a type of SPPA whereby the positions of points remains the same, but the labels (here, fragment versus whole specimen) are randomly and repeatedly permuted (Jacquemyn et al., 2010; Mitchell et al., 2018; Raventós et al., 2010). As such, RLAs do not measure spatial segregation or aggregation and therefore do not test processes that resulted in the fossil’s location, but instead measure the differences in the spatial distributions of the permuted labels (Getzin et al., 2008). In order to test if fragments appeared in areas of high *Fractofusus* densities (high density-dependence between fragments and whole specimens), we used the difference test which detects where one type of pattern is in areas of high density of the joined point pattern (Wiegand & Moloney, 2014), and so this test was used to detect density-dependence of the fragments in the *Fractofusus andersoni* population on Brasier surface in Programita (v November 2018) (Wiegand, 2018).

Diggle’s goodness-of-fit test (*p_d_*) was then used to quantitatively assess differences between the observed and simulated PCFs generated by univariate SPPA and RLA. Diggle’s goodness-of-fit test provides a hypothesis test, with the alternative hypothesis being that the measured process (here, PCF of the fossil’s point pattern) departs from the modelled process over a specified, biologically-relevant distance interval (e.g., 2 – 10 cm) (Diggle, 2003). A high *p_d_* is interpreted to be a good model fit, alongside visual inspection of the simulation envelopes of the PCF plot and a low *p_d_* interpreted to be a bad model fit (Diggle, 2003; Wiegand & Moloney, 2014).

### Identifying outliers & impacts on population and community ecology

Rosner tests enable the quantitative identification of multiple outliers from a distribution of data. Rosner tests produce a test statistic, *R,*

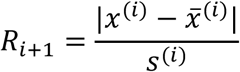

where 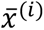 is the sample mean and *s*^(*i*)^ is the standard deviation after the *i* most extreme observations have been removed. The hypothesis that there are *k* outliers is tested by comparing *R_k_* to the critical value *λ_k_* for a significance level *α* (here, *α* = 0.05) with outliers present where *R_k_* > *λ_k_* (Millard, 2013). For the R value, the larger the R value, the greater the outlier is from the rest of the distribution.

We used Rosner tests from the EnvStats package (Millard, 2013) in R on the height distribution of all abundant (n > 30) populations in our dataset to identify outliers of height distributions inferred to be potential survivors. We used height for all upright taxa, modelling height for non-upright taxa *Fractofusus* and *Hapsidophyllas* as one third of the frond width *cf* (Mitchell & Kenchington, 2018). Statistically significant outliers were then assessed visually and substantial outliers (> 10 cm difference from the population mean height) were considered different enough to be survivors.

Since, by definition, outliers are few individuals, we had a low sample size with which to investigate their impact on population dynamics. We therefore noted whether outliers were representative of the most abundant taxa on a given surface. Kolmogorov-Smirnov (KS) tests (Kolmogorov, 1933; Smirnov, 1948) were used to test if there were systematic changes in all population densities on a given surface with distance from each outlier with a heterogenous Poisson distribution compared to what was expected by random. KS tests were performed in the spatstat package in R (Baddeley, 2015, 2024).

## Results

### Fragments

We identified 407 complete *Fractofusus andersoni* specimens and 28 fragment specimens (Fig. 1) on the Brasier surface (Fig. 2). We found that fragments had a significantly larger width than complete *Fractofusus* specimens (mean frond width: complete specimens = 1.05 cm, fragment specimens = 3.40 cm; W = 1,737, *p* < 0.001; Fig. 3).

**Figure 3:**
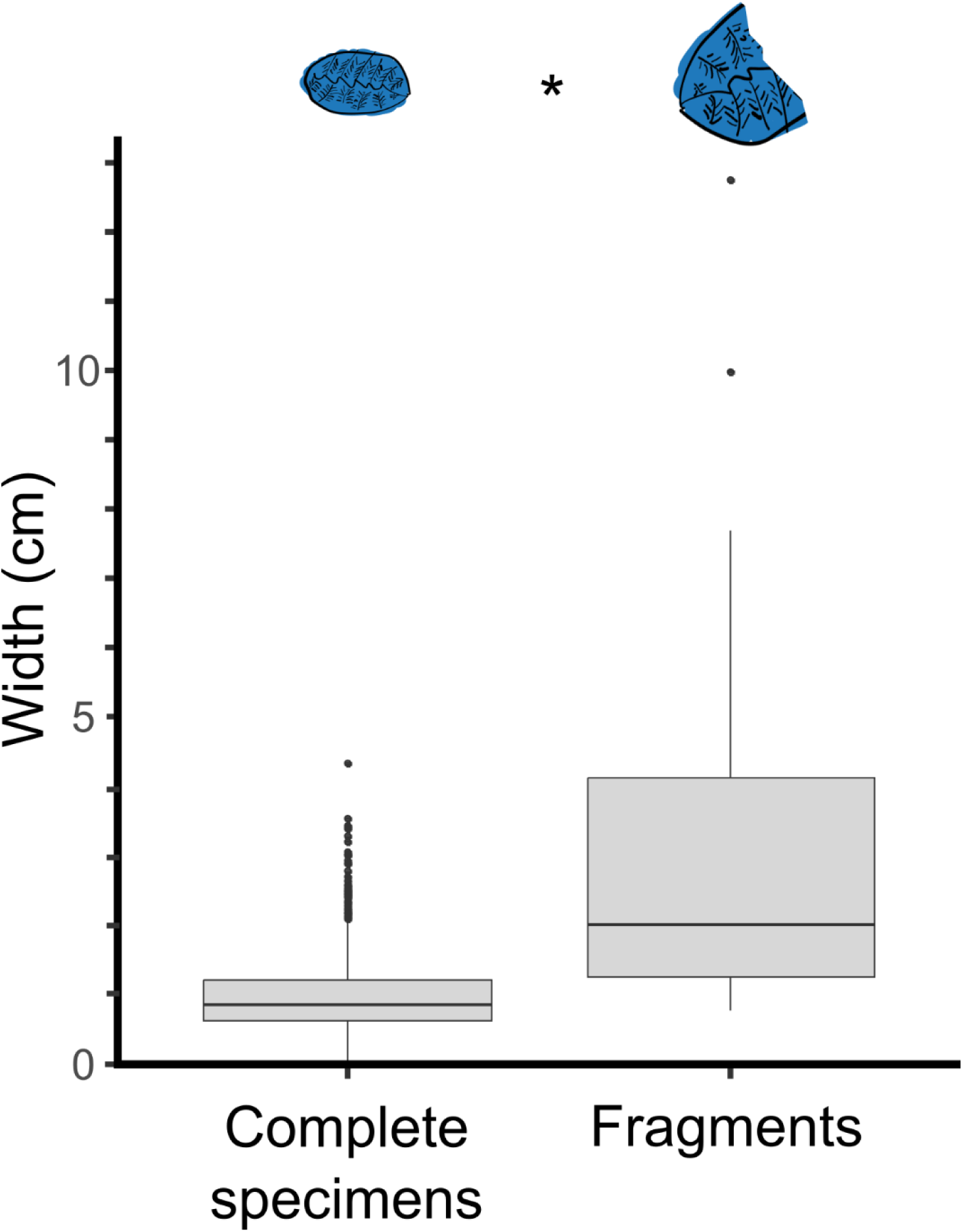
Differences in the width of fragments and complete specimens of *Fractofusus andersoni* on the Brasier surface. Asterisk denotes a statistically significant difference.

Complete *Fractofusus andersoni* specimens on the Brasier surface were clustered in areas where fragment density was highest, indicated by a good fit to an ITC model (*p_d_* = 0.793, range = 0 – 20 cm; Fig. 4a), indicating reproductive clusters forming in high fragment-density areas. In comparison, a CSR model (*p_d_* < 0.001, range = 0 – 20 cm) indicating no biotic or abiotic effects, a TC model (*p_d_* < 0.001, range = 0 – 20 cm) indicating independence from the fragments, and a HP model (*p_d_* = 0.004, range = 0 – 20 cm) indicating no clustering within areas of high habitat preference all had poor fit.

**Figure 4:**
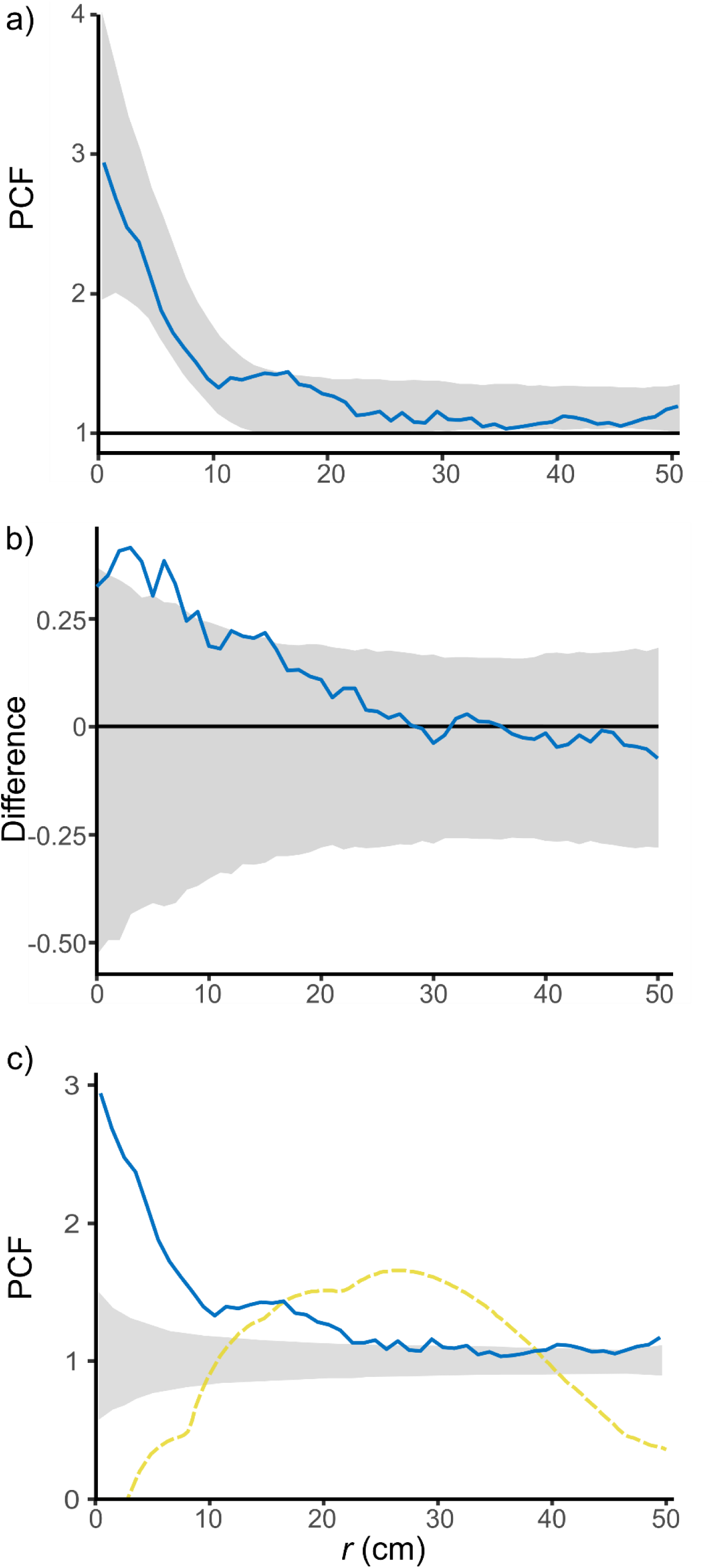
a) Pair correlation function (PCF) plot for inhomogenous Thomas cluster model for *Fractofusus andersoni* on Brasier surface with the inhomogenous background generated from the density of fragments. b) PCF plot for density-dependent random labelling of fragments on *Fractofusus andersoni* on the Brasier surface. PCF in blue, simulation envelope in grey. Excursions above the simulation envelope indicates more fragments are present in areas of higher *Fractofusus* densities at a given distance *r*. c) PCFs of complete (blue) and fragmented (yellow) *Fractofusus andersoni* specimens on the Brasier surface. Envelope generated from complete spatial randomness model of complete *Fractofusus* specimens.

We found that fragments were in areas of high *Fractofusus* densities indicated by a deviation from CSR using RLA at small spatial scales (*p_d_* = 0.059, range = 2 – 10 cm; Fig. 4b).

The RLA PCFs shows us that the fragments and whole *Fractofusus* on the Brasier surface are not alike, which suggests that the spatial patterns are unlikely to be caused by similar ecological processes (Fig. 4c).

### Survivors

Eleven outlier specimens were identified from Rosner Tests and subsequent visual assessment from seven populations across four surfaces (Table 1; Table S1): three *Charnia*, two *Primocandelabrum* spp., and one *Charniodiscus* sp. specimens on Bed B; two “Taxon B” (an undescribed unifoliate rangeomorph frond) specimens on Brasier surface; one *Bradgatia* sp. on the Bishop’s Cove surface; and one *Charniodiscus procerus* specimen on the E Surface (Fig. 5). An outsized *Primocandelabrum* sp. specimen on H26 was also identified qualitatively, but the population on H26 was too small to apply subsequent quantitative analyses (Table S1). All of these outlier specimens were therefore interpreted to be survivors from a disturbance event *cf* (Wilby et al., 2015). We observed that in nine of eleven cases, the survivors represented the most abundant taxa on their respective surfaces, with the exception of “Taxon B”, which was second most abundant behind *Fractofusus andersoni* on the Brasier surface, and *Charniodiscus procerus* on the E surface (Table S2). Notably, across all seven populations, there was no apparent signal from the presence or absence of stems: *Primocandelabrum* and *Charniodiscus* have stems and *Charnia, Bradgatia*, and “Taxon B” do not have stems.

**Figure 5:**
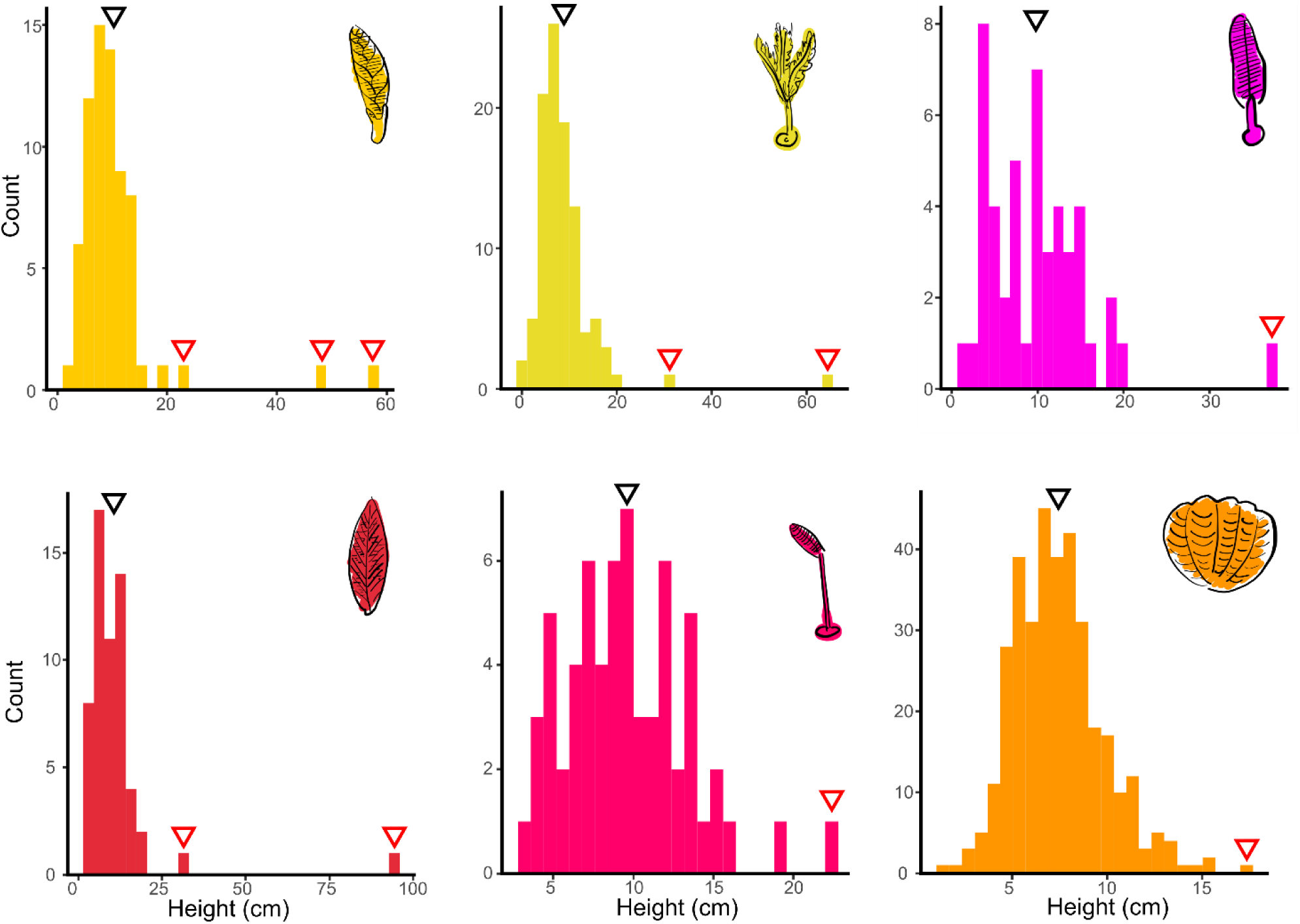
Height distributions of the outlier specimens (red arrows) identified by Rosner Tests with > 10 cm difference from the population’s mean height (black arrows). Cartoons represent taxa right to left top row: yellow *Charnia* on Bed B; yellow *Primocandelabrum* on Bed B; pink *Charniodiscus* on Bed B; bottom row: red “Taxon B” on Brasier surface; pink *Charniodiscus procerus* on E Surface; orange *Bradgatia* sp. on Bishop’s Cove.

**Table 1:** Populations with outliers identified by Rosner Tests. Each Rosner statistic (*R*_1_, *R*_2_, and *R*_3_) represents the statistic for an outsized individual.

| Surface | Taxon | $R_1$ | $R_2$ | $R_3$ |
| --- | --- | --- | --- | --- |
| Brasier | "Taxon B" | 6.78 | 4.25 |  |
| Bed B | *Charnia masoni* | 5.50 | 6.70 | 3.38 |
| Bed B | *Charniodiscus* spp. | 4.49 |  |  |
| Bed B | *Primocandelabrum* spp. | 7.70 | 4.86 |  |
| H26 | *Primocandelabrum* sp. | 3.52 |  |  |
| Bishop's Cove | *Bradgatia* sp. | 4.18 |  |  |
| E Surface | *Charniodiscus procerus* | 3.37 |  |  |

All KS tests comparing a heterogenous Poisson distributed intensity from each outlier to a random distribution were non-significant (i.e., *p* > 0.05) (Table S3), indicating no association between outsize specimens (survivors) and conspecific population densities.

## Discussion

After environmental disturbance, extant organisms use both short- and long-distance dispersal processes to recolonise environments (Brunner et al., 2022; Burt et al., 2024; Kim et al., 2022; Wild et al., 2014). We tested how two types of local recolonisation processes – reproductively-active fragments of *Fractofusus* and outsize (presumed surviving) individuals – influenced the post-disturbance communities in the Avalon. Our spatial analyses showed that fragments of *Fractofusus* on the Brasier surface showed a positive density-dependent correlation with smaller, complete *Fractofusus* fossils. The dominance of *Fractofusus* on this surface (77.1% relative abundance), coupled with our results, are consistent with the hypothesis that fragments were reproductively active on the Brasier surface and aided in recolonisation and therefore influenced secondary community succession. Our results provide evidence that fragmentary reproduction is an effective re-colonisation mechanism (Fig. 5) and provide the first evidence supporting reproduction from fragments (as opposed to putative budding, water-borne propagules, or stolon) in early animals in the Avalon. It is noteworthy that *Fractofusus* appears to be the most reproductively plastic Avalon taxon (water-borne (Mitchell et al., 2015, Darroch et al. 2013), stolon (Delahooke et al., 2024; Liu & Dunn, 2020; Mitchell et al., 2015), and fragmentary reproduction described here), although detection of reproductive plasticity could be due to high abundance of *Fractofusus* enabling a higher likelihood of detection. The Brasier surface is an early succession community (Stephenson et al., 2024), so, because Avalon communities do not undergo turnover during succession but instead accumulate diversity while maintaining similar community compositions, it is plausible that the Brasier community could have matured to look similar to later-stage, *Fractofusus andersoni-*dominated communities such as the Goldmine or H14 surfaces. On the D, E, and H14 surfaces – all dominated by *Fractofusus*, but later in succession than Brasier (Stephenson et al., 2024) – we observed *Fractofusus* fragments, but in much lower relative abundances (0.1%, 0.3%, and 0.3% on D, E, and H14 surfaces, respectively; Table S2). Fragments on later-stage surfaces may indicate that fragmentation could have influenced the community ecology of these surfaces earlier in succession alongside presumed long-distance, water-borne propagule effects (Clapham et al., 2003; Eden et al., 2022; Mitchell et al., 2015) by contributing to *Fractofusus* dominance. These cryptic fragments could result in “fairy ring”-like patterns of *Fractofusus andersoni*, which could be areas where a fragment was present, but has since decayed – a pattern observed on the H14 surface (Fig. 6).

**Figure 5:**
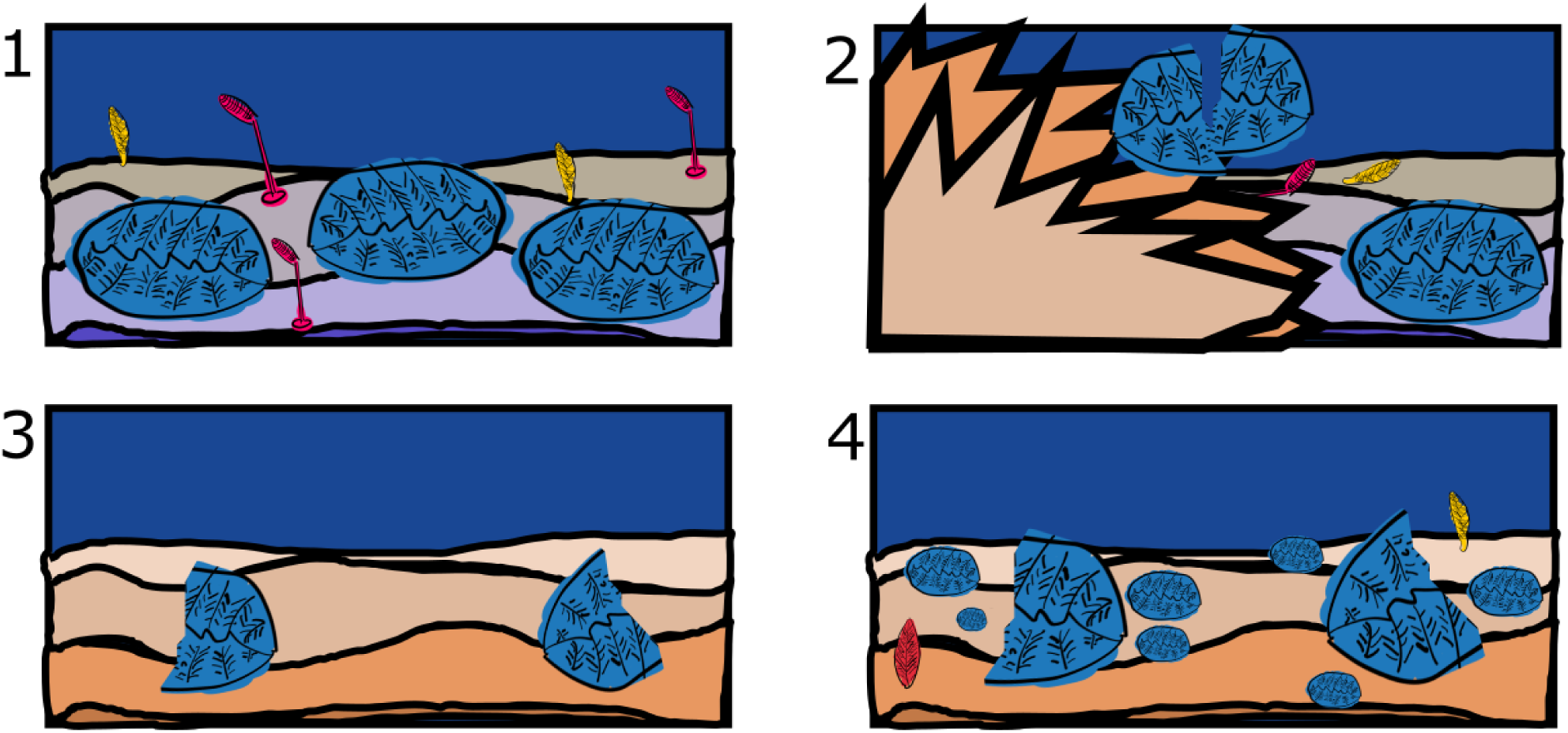
Schematic of the processes leading to secondary *Fractofusus* colonisation on the Brasier surface. Panel 1: Later-succession, pre-disturbance *Fractofusus*-dominated community; Panel 2: A sedimentary disturbance; Panel 3: Fragmentary remains of *Fractofusus* (blue) generated from the pre-disturbance community settled onto the post-disturbance substrate; Panel 4: the post-disturbance community later in succession – secondary processes have led to the dominance of *Fractofusus* due to the ecological impacts of fragments, and other taxa (yellow and red frondomorphs) have arrived via long-distance dispersal but have lower abundances.

**Figure 6:**
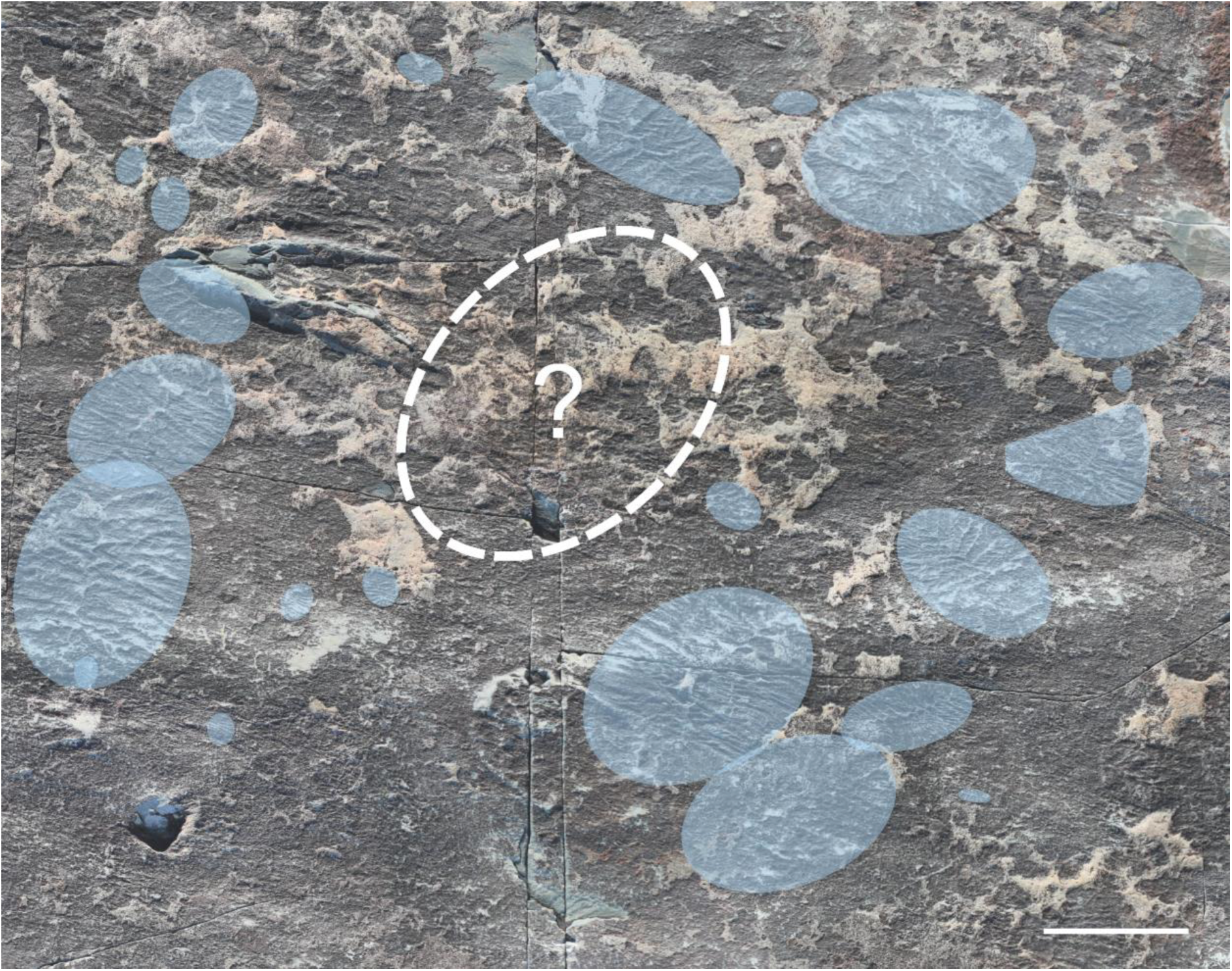
A “fairy ring”-like structure of *Fractofusus andersoni* on the H14 surface in Discovery Geopark, Newfoundland, Canada. Blue ellipses indicate *Fractofusus andersoni* fossils, and dashed outline where a possible fragment could have once been but has since decayed away before preservation of the H14 community. Scale bar ∼ 5 cm.

In contrast to the fragments, surviving fronds (those that were an outlier in a population’s height distribution) had minimal impact on the spatial population ecology in the eleven instances in which they were found. However, we note that, like (Wilby et al., 2015), nine of eleven outlier individuals were representative of one of the most common taxa where they were found (the exceptions being “Taxon B” on the Brasier surface, which was the second most abundant taxa (10.9% relative abundance), and *Charniodiscus procerus* on the E Surface (2.9% relative abundance)). While not included in the mapped areas of our dataset, we are aware of at least two other instances of outlier individuals that do not represent the most abundant taxon in the community: the holotype *Pectinifrons* specimen on Mistaken Point North (Figure 4 in (Bamforth et al., 2008)) is much larger than all other specimens on the surface which also have low abundance; and the large *Frondophyllas grandis* specimen on Lower Mistaken Point (Figure 2.2 in (Bamforth & Narbonne, 2009)) which is one of only three *Frondophyllas* specimens on the surface. Our results show that there is no significant benefit to survival of disturbances through increased height, suggesting that height may not have evolved to survive disturbance events.

Our results show that outlier specimens are present in four of 19 communities, with outliers representing the most abundant taxa for five of seven populations (on two out of four communities). The taxonomic identity of outliers results from a combination of random chance (which specimens get smothered, or not), and the relative abundances of the species (the greater the abundance, the more likely a single representative is to survive). The variation of taxonomic identity (Fig. 5), shows a lack of species-specific signal to survival. Therefore, the outliers were likely to be from the most abundant taxa within the pre-disturbance communities. The similarities between the taxonomic identification of the outliers with the most abundant taxa in the post-disturbance communities, suggest that the sources for both pre- and post-disturbance colonisations were similar and, because our results (Table S3) excluded local-recolonisation from the outlier individuals (i.e., there is no increased density around or downstream of the outliers of conspecifics), instead these results suggest that it is long-distance dispersal (i.e., from outside the mapped communities) that is seeding new communities. Our results instead reinforce the role of height in reproduction for upright Avalon taxa since increased height increases dispersal distance, but not local survival (Gaylord et al., 2002; Mitchell & Kenchington, 2018), and provides further evidence of the importance of long-distance dispersal processes to Avalon community ecology (Mitchell et al., 2019, 2020; Mitchell & Kenchington, 2018).

The impact of long-distance dispersal and colonisation has been found through the primary colonisation of barren substrates by waterborne *Fractofusus* (Mitchell et al. 2015), with lasting effects through subsequent development because species turnover, environmental filtering, and interspecific interactions throughout succession are limited (Stephenson et al., 2024). Between community (within metacommunity) evidence of the importance of dispersal limitation is seen in the metacommunity structure of the Avalon, where species-poor communities are subsets of species-rich communities (Eden et al., 2022). In this structure, species-poor communities correspond to species-specific characteristics, which is a pattern found where there is a large species pool, with long-dispersal patterns leading to different community compositions (Presley et al., 2010). However, once the substrate has been colonised, local reproductive processes (fragmentation and different stoloniferous strategies (Delahooke et al., 2024; Liu & Dunn, 2020; Mitchell et al., 2015), and putative budding (Pasinetti & McIlroy, 2023)) appear to have strong impact on subsequent community succession. The capacity of Avalon taxa to exhibit both long-distance (Darroch et al. 2013, Mitchell et al. 2015) and these local reproductive modes (our results; (Delahooke et al., 2024; Mitchell et al., 2015)), perhaps mediated by the intensity of environmental disturbance, could be key in dictating community composition.

In modern, regularly-disturbed ecosystems, biological legacies are important contributors to post-disturbance community ecology (Bače et al., 2015; Nyström & Folke, 2001; Pérez-Hernández & Gavilán, 2021; Wild et al., 2014). In many modern deep-sea environments, however, there is an emphasis on long-distance metacommunity structures, particularly in isolated environments such as hydrothermal vents where long-distance dispersal is an important component of community structuring (Brunner et al., 2022). Similarly, patchy disturbance generated by sedimentation events (such as turbidites in the Avalon) is likely to have produced large-scale heterogeneity and mosaic effects (Bigham et al., 2021), with long-distance dispersal helping to populate these patches. Given a global distribution of taxa such as *Charnia* in the Ediacaran (Boddy et al., 2022; Darroch et al., 2013; Grazhdankin et al., 2008; Wu et al., 2024), and an Avalon metacommunity structure whereby the most diverse sites reflect almost all the known diversity (Eden et al., 2022), coupled with our results showing the importance of local colonisation strategies, it is likely that similar long-distance adaptations were in place in the Avalon. Similar patterns of intense disturbance alongside dispersal limitations are seen in the Antarctic deep sea (Griffiths et al., 2023) where most sessile organisms are dispersal-limited (Thatje, 2012) shaping high beta diversity (Thrush et al., 2010). Disturbance in the form of ice scours provides the major form of ecological turnover (Thatje, 2012) in the absence of widespread predation (Khan et al., 2024; Smith et al., 2017), so that the survival of disturbance and occupation of refugia is of substantial ecological importance (Barnes & Conlan, 2007; Potthoff et al., 2006). Therefore, the importance of a combination of long-distance dispersal and local re-colonisation effects in the Avalon is strikingly similar to trade-offs in modern deep-sea organisms, as alternative colonisation strategies in the Avalon reflect a combination of disturbance responses alongside the influence of long-distance dispersal from the surrounding metacommunity.

## Acknowledgements

Funding was provided by a School of Biological Sciences Balfour Studentship (PFAG/076) to NPS; a Natural Environment Research Council C-CLEAR DTP studentship (LBAG/265.03.G10) to KMD; a Natural Environment Research Council C-CLEAR DTP studentship (NE/S007164/1, with project reference 2889755) to PAB; Leverhulme Trust Funding (ECF-2018-542) and from the Isaac Newton Trust (INT18.08(h)) to CGK; a Natural Environment Research Council Standard Grant (NE/P002412/1) and an Independent Research Fellowship (NE/S014756/1) to EGM. The Parks and Natural Areas Division, Department of Environment and Conservation, Government of Newfoundland and Labrador, provided permits to conduct research within the Mistaken Point Ecological Reserve (MPER) in 2010, 2016 - 2019, and 2021 - 2023. Readers are advised that access to MPER is by scientific research permit only. Fossil surfaces in the Bonavista Peninsula and Bay Roberts area are protected under Reg. 67/11, of the Historic Resources Act 2011 and their access is only allowed under permit from the Government of Newfoundland and Labrador. Fieldwork in the Discovery Geopark and the Bay Roberts area was conducted under permit from the Government of Newfoundland and Labrador. Bed B forms part of a designated Site of Special Scientific Interest (SSSI), protected in law and administered by Natural England. Access to Bed B casts was facilitated by BGS and P. Wilby. We thank F. Dunn for helpful discussion. We thank E. Samson for everything she does to support us and our work.

## Author contributions

This work was conceptualised by NPS and EGM. Data collection was carried out by NPS, KMD, NB, BWTR, CGK, and EGM, and the data were processed by NPS, KMD, and EGM. Statistical analyses were conducted by NPS, KMD, and EGM. The original draft was written by NPS, PAB, AM, and EGM, and all authors contributed to the review and editing of the final manuscript.

## Supplementary material

**Table S1:**
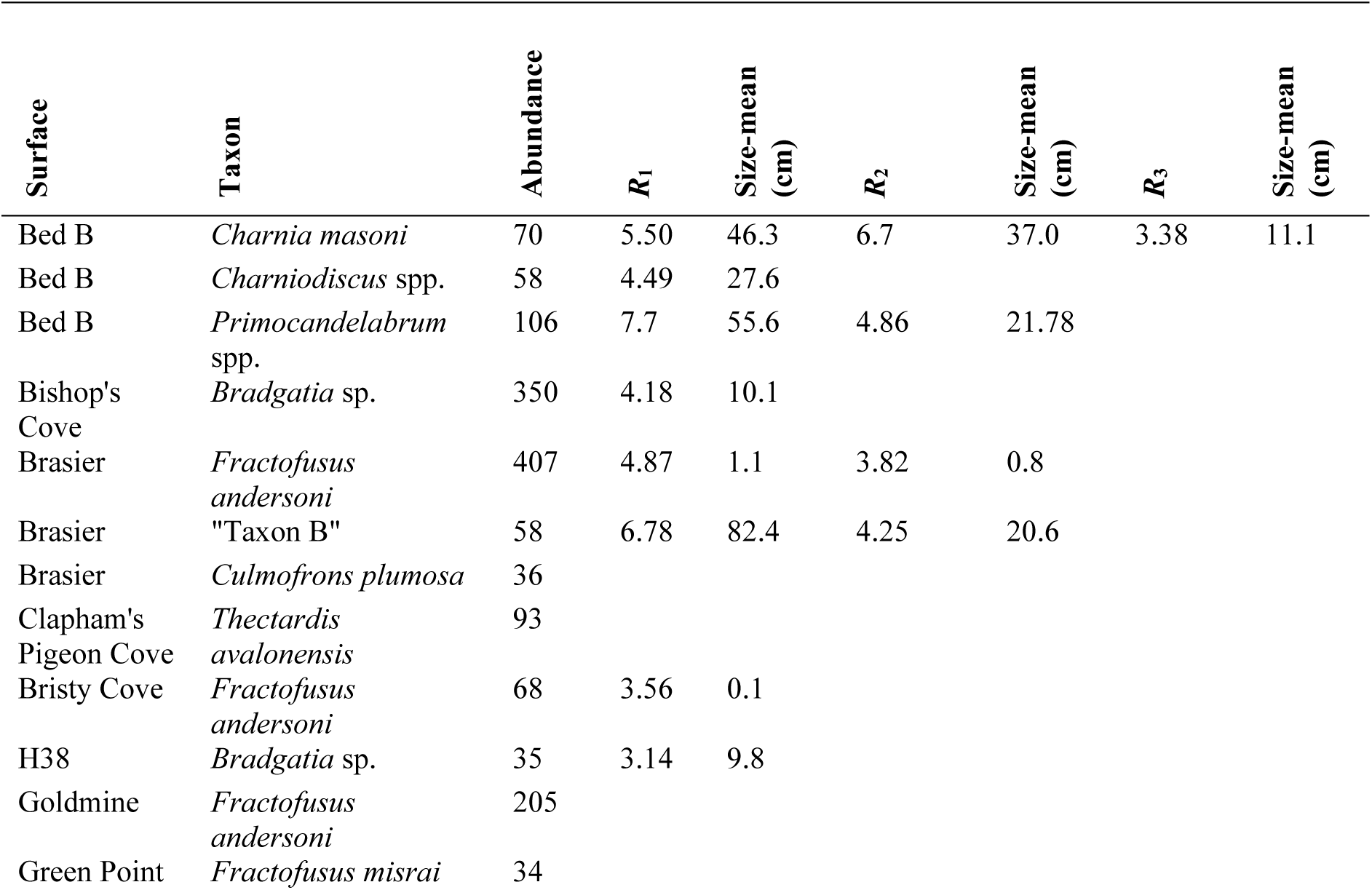
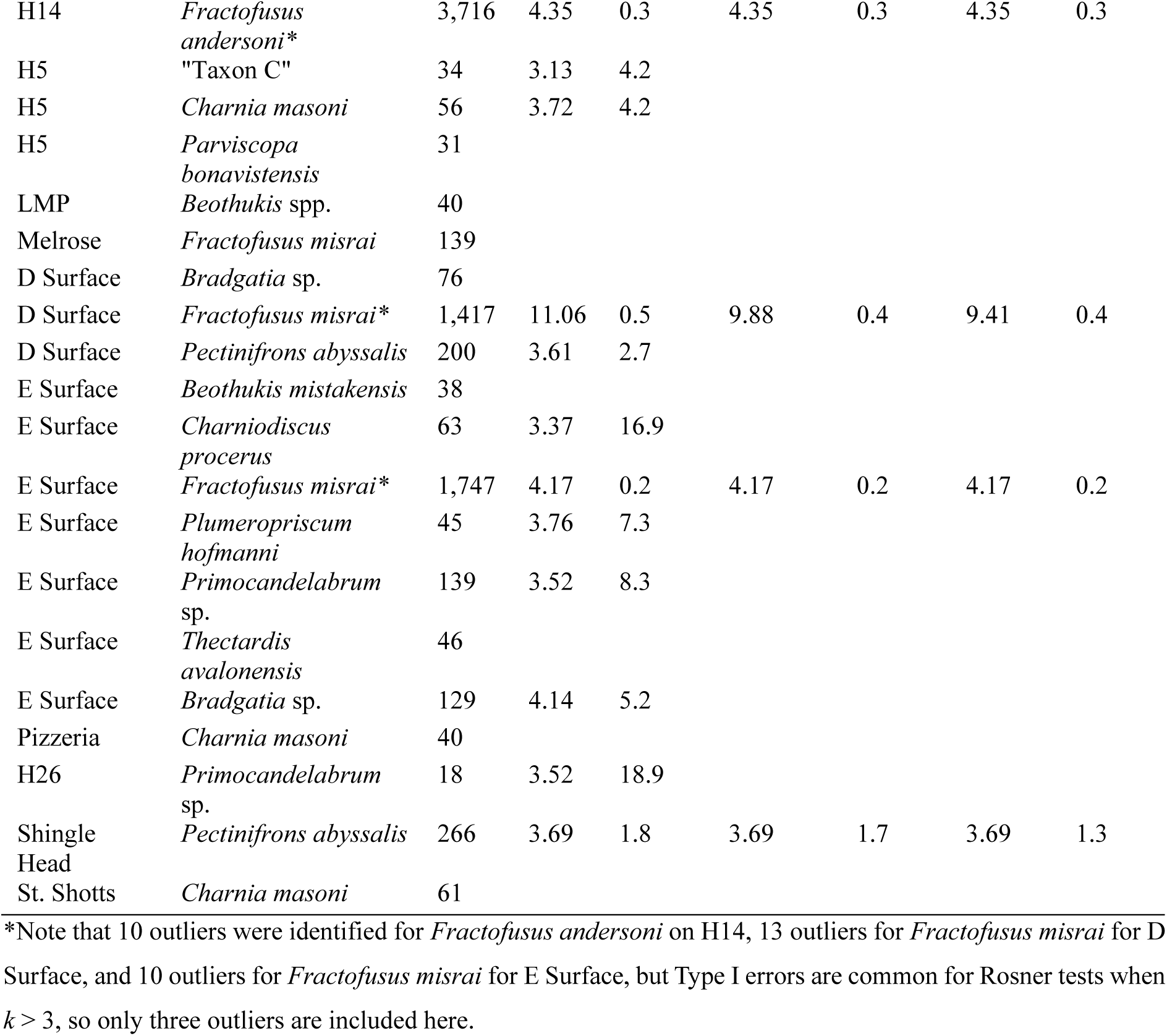
Rosner test results for all abundant (n > 30) populations. Size-mean columns indicate the difference between the size (height, length, or width) of the outlier and the population mean.

| Surface | Taxon | Abundance | $R_1$ | Size-mean<br>(cm) | $R_2$ | Size-mean<br>(cm) | $R_3$ | Size-mean<br>(cm) |
| --- | --- | --- | --- | --- | --- | --- | --- | --- |
| Bed B | *Charnia masoni* | 70 | 5.50 | 46.3 | 6.7 | 37.0 | 3.38 | 11.1 |
| Bed B | *Charniodiscus* spp. | 58 | 4.49 | 27.6 |  |  |  |  |
| Bed B | *Primocandelabrum*<br>spp. | 106 | 7.7 | 55.6 | 4.86 | 21.78 |  |  |
| Bishop's<br>Cove | *Bradgatia* sp. | 350 | 4.18 | 10.1 |  |  |  |  |
| Brasier | *Fractofusus*<br>*andersoni* | 407 | 4.87 | 1.1 | 3.82 | 0.8 |  |  |
| Brasier | "Taxon B" | 58 | 6.78 | 82.4 | 4.25 | 20.6 |  |  |
| Brasier | *Culmofrons plumosa* | 36 |  |  |  |  |  |  |
| Clapham's<br>Pigeon Cove | *Thectardis*<br>*avalonensis* | 93 |  |  |  |  |  |  |
| Bristy Cove | *Fractofusus*<br>*andersoni* | 68 | 3.56 | 0.1 |  |  |  |  |
| H38 | *Bradgatia* sp. | 35 | 3.14 | 9.8 |  |  |  |  |
| Goldmine | *Fractofusus*<br>*andersoni* | 205 |  |  |  |  |  |  |
| Green Point | *Fractofusus misrai* | 34 |  |  |  |  |  |  |
| H14 | *Fractofusus andersoni* * | 3,716 | 4.35 | 0.3 | 4.35 | 0.3 | 4.35 | 0.3 |
| H5 | "Taxon C" | 34 | 3.13 | 4.2 |  |  |  |  |
| H5 | *Charnia masoni* | 56 | 3.72 | 4.2 |  |  |  |  |
| H5 | *Parviscopa bonavistensis* | 31 |  |  |  |  |  |  |
| LMP | *Beothukis* spp. | 40 |  |  |  |  |  |  |
| Melrose | *Fractofusus misrai* | 139 |  |  |  |  |  |  |
| D Surface | *Bradgatia* sp. | 76 |  |  |  |  |  |  |
| D Surface | *Fractofusus misrai* * | 1,417 | 11.06 | 0.5 | 9.88 | 0.4 | 9.41 | 0.4 |
| D Surface | *Pectinifrons abyssalis* | 200 | 3.61 | 2.7 |  |  |  |  |
| E Surface | *Beothukis mistakensis* | 38 |  |  |  |  |  |  |
| E Surface | *Charniodiscus procerus* | 63 | 3.37 | 16.9 |  |  |  |  |
| E Surface | *Fractofusus misrai* * | 1,747 | 4.17 | 0.2 | 4.17 | 0.2 | 4.17 | 0.2 |
| E Surface | *Plumeropriscum hofmanni* | 45 | 3.76 | 7.3 |  |  |  |  |
| E Surface | *Primocandelabrum* sp. | 139 | 3.52 | 8.3 |  |  |  |  |
| E Surface | *Thectardis avalonensis* | 46 |  |  |  |  |  |  |
| E Surface | *Bradgatia* sp. | 129 | 4.14 | 5.2 |  |  |  |  |
| Pizzeria | *Charnia masoni* | 40 |  |  |  |  |  |  |
| H26 | *Primocandelabrum* sp. | 18 | 3.52 | 18.9 |  |  |  |  |
| Shingle Head | *Pectinifrons abyssalis* | 266 | 3.69 | 1.8 | 3.69 | 1.7 | 3.69 | 1.3 |
| St. Shotts | *Charnia masoni* | 61 |  |  |  |  |  |  |

\*Note that 10 outliers were identified for *Fractofusus andersoni* on H14, 13 outliers for *Fractofusus misrai* for D Surface, and 10 outliers for *Fractofusus misrai* for E Surface, but Type I errors are common for Rosner tests when $k > 3$ , so only three outliers are included here.

**Table S2:** Abundances for all taxa across 19 surfaces.

| Taxon | Bed B | Bishop's Cove | Brasier | Bristy Cove | Clapham's Pigeon Cove | H38 | Goldmine | Green Point | H5 | H14 | H26 | LMP | Melrose | D Surface | E Surface | G Surface | Pizzeria | Shingle Head | St. Shotts |
| --- | --- | --- | --- | --- | --- | --- | --- | --- | --- | --- | --- | --- | --- | --- | --- | --- | --- | --- | --- |
| *Arborea arborea* | 15 | 0 | 0 | 0 | 0 | 0 | 0 | 0 | 0 | 0 | 0 | 1 | 0 | 0 | 0 | 0 | 0 | 0 | 0 |
| *Auroralumina attenboroughii* | 1 | 0 | 0 | 0 | 0 | 0 | 0 | 0 | 0 | 0 | 0 | 0 | 0 | 0 | 0 | 0 | 0 | 0 | 0 |
| *Bradgatia linfordensis* | 9 | 0 | 0 | 0 | 0 | 0 | 0 | 0 | 0 | 0 | 0 | 0 | 0 | 0 | 0 | 0 | 0 | 0 | 0 |
| "Brush" | 11 | 0 | 0 | 0 | 0 | 0 | 0 | 0 | 0 | 0 | 0 | 0 | 0 | 0 | 0 | 0 | 0 | 0 | 0 |
| *Charnia masoni* | 70 | 3 | 6 | 0 | 0 | 0 | 0 | 0 | 56 | 2 | 7 | 11 | 0 | 2 | 12 | 0 | 40 | 0 | 61 |
| *Charniodiscus* spp. | 48 | 0 | 0 | 0 | 0 | 3 | 0 | 0 | 0 | 0 | 0 | 0 | 0 | 0 | 7 | 0 | 3 | 1 | 2 |
| *Hiemalora stellaris* | 2 | 11 | 8 | 1 | 0 | 1 | 3 | 1 | 2 | 4 | 3 | 3 | 1 | 4 | 23 | 0 | 4 | 0 | 0 |
| *Hyllaeacullus fordi* | 12 | 0 | 0 | 0 | 0 | 0 | 0 | 0 | 0 | 0 | 0 | 0 | 0 | 0 | 0 | 0 | 0 | 0 | 0 |
| "Taxon A" | 5 | 0 | 2 | 0 | 0 | 0 | 0 | 0 | 0 | 1 | 0 | 0 | 0 | 0 | 0 | 0 | 1 | 0 | 0 |
| *Primocandelabrum aethelflaedia* | 5 | 0 | 0 | 0 | 0 | 0 | 0 | 0 | 0 | 0 | 0 | 0 | 0 | 0 | 0 | 0 | 0 | 0 | 0 |
| *Primocandelabrum boyntoni* | 9 | 0 | 0 | 0 | 0 | 0 | 0 | 0 | 0 | 0 | 0 | 0 | 0 | 0 | 0 | 0 | 0 | 0 | 0 |
| *Primocandelabrum* sp. | 92 | 1 | 8 | 0 | 0 | 5 | 2 | 1 | 8 | 2 | 18 | 22 | 0 | 6 | 139 | 8 | 10 | 0 | 4 |
| *Bradgatia* sp. | 0 | 350 | 0 | 0 | 0 | 35 | 0 | 8 | 1 | 0 | 0 | 0 | 11 | 76 | 129 | 28 | 1 | 13 | 0 |
| *Trepassia wardae* | 0 | 1 | 0 | 0 | 0 | 0 | 0 | 0 | 0 | 0 | 0 | 1 | 0 | 0 | 1 | 0 | 0 | 0 | 0 |
| *Charniodiscus arboreus* | 0 | 0 | 11 | 0 | 0 | 1 | 2 | 2 | 26 | 1 | 1 | 0 | 0 | 0 | 0 | 0 | 8 | 0 | 2 |
| *Culmofrons plumosa* | 0 | 0 | 36 | 1 | 0 | 0 | 0 | 0 | 0 | 0 | 0 | 6 | 0 | 0 | 5 | 1 | 0 | 0 | 0 |
| "Taxon B" | 0 | 0 | 58 | 0 | 0 | 0 | 0 | 0 | 0 | 0 | 0 | 0 | 0 | 0 | 0 | 0 | 0 | 0 | 0 |
| *Fractofusus andersoni* | 0 | 0 | 407 | 68 | 0 | 1 | 205 | 0 | 6 | 3,716 | 2 | 8 | 0 | 0 | 0 | 0 | 0 | 0 | 0 |
| *Fractofusus* fragment | 0 | 0 | 28 | 0 | 0 | 0 | 0 | 0 | 0 | 13 | 0 | 0 | 0 | 4 | 18 | 0 | 0 | 0 | 0 |
| *Plumeropriscum hofmanni* | 0 | 0 | 1 | 0 | 0 | 0 | 0 | 0 | 0 | 0 | 0 | 4 | 0 | 0 | 45 | 0 | 0 | 0 | 0 |
| Lobate disc | 0 | 0 | 0 | 0 | 1 | 0 | 0 | 0 | 0 | 0 | 0 | 1 | 0 | 2 | 70 | 0 | 4 | 0 | 0 |
| *Thectardis avalonensis* | 0 | 0 | 0 | 0 | 93 | 0 | 1 | 0 | 0 | 2 | 0 | 0 | 0 | 0 | 46 | 2 | 2 | 0 | 0 |
| *Vinlandia antecedens* | 0 | 0 | 0 | 0 | 4 | 0 | 0 | 0 | 0 | 0 | 0 | 1 | 0 | 0 | 0 | 0 | 0 | 0 | 0 |
| "Taxon E" | 0 | 0 | 0 | 0 | 0 | 2 | 1 | 1 | 0 | 0 | 0 | 10 | 0 | 6 | 3 | 0 | 0 | 0 | 0 |
| *Charniodiscus spinosus* | 0 | 0 | 0 | 0 | 0 | 2 | 0 | 0 | 0 | 0 | 0 | 11 | 0 | 1 | 24 | 0 | 4 | 0 | 0 |
| *Fractofusus misrai* | 0 | 0 | 0 | 0 | 0 | 0 | 0 | 34 | 0 | 0 | 0 | 0 | 139 | 1,417 | 1,747 | 0 | 0 | 2 | 0 |
| *Pectinifrons abyssalis* | 0 | 0 | 0 | 0 | 0 | 0 | 0 | 9 | 0 | 4 | 0 | 0 | 24 | 200 | 0 | 0 | 0 | 266 | 0 |
| "Taxon C" | 0 | 0 | 0 | 0 | 0 | 0 | 0 | 0 | 34 | 0 | 0 | 22 | 0 | 0 | 0 | 0 | 0 | 0 | 0 |
| *Parviscopa bonavistensis* | 0 | 0 | 0 | 0 | 0 | 0 | 0 | 0 | 31 | 0 | 0 | 0 | 0 | 0 | 0 | 0 | 0 | 0 | 0 |
| *Hadryniscala avalonica* | 0 | 0 | 0 | 0 | 0 | 0 | 0 | 0 | 0 | 1 | 0 | 0 | 0 | 0 | 0 | 0 | 0 | 0 | 0 |
| "Taxon D" | 0 | 0 | 0 | 0 | 0 | 0 | 0 | 0 | 0 | 4 | 0 | 0 | 0 | 0 | 0 | 0 | 0 | 0 | 0 |
| *Beothukis mistakensis* | 0 | 0 | 0 | 0 | 0 | 0 | 0 | 0 | 0 | 0 | 0 | 8 | 0 | 0 | 38 | 2 | 0 | 0 | 0 |
| *Fronodophyllas grandis* | 0 | 0 | 0 | 0 | 0 | 0 | 0 | 0 | 0 | 0 | 0 | 2 | 0 | 0 | 0 | 0 | 4 | 0 | 0 |
| "Ostrich feather" | 0 | 0 | 0 | 0 | 0 | 0 | 0 | 0 | 0 | 0 | 0 | 15 | 0 | 0 | 0 | 0 | 0 | 0 | 0 |
| "Taxon F" | 0 | 0 | 0 | 0 | 0 | 0 | 0 | 0 | 0 | 0 | 0 | 0 | 0 | 1 | 23 | 0 | 0 | 0 | 0 |
| *Hapsidophyllas flexibilis* | 0 | 0 | 0 | 0 | 0 | 0 | 0 | 0 | 0 | 0 | 0 | 0 | 0 | 1 | 1 | 0 | 0 | 0 | 0 |
| "Taxon G" | 0 | 0 | 0 | 0 | 0 | 0 | 0 | 0 | 0 | 0 | 0 | 0 | 0 | 0 | 11 | 0 | 0 | 0 | 0 |
| *Charniodiscus procerus* | 0 | 0 | 0 | 0 | 0 | 0 | 0 | 0 | 0 | 0 | 0 | 0 | 0 | 0 | 63 | 8 | 6 | 0 | 0 |
| *Charniodiscus concentricus* | 0 | 0 | 0 | 0 | 0 | 0 | 0 | 0 | 0 | 0 | 0 | 0 | 0 | 0 | 0 | 0 | 1 | 0 | 0 |
| "Taxon H" | 0 | 0 | 0 | 0 | 0 | 0 | 0 | 0 | 0 | 0 | 0 | 0 | 0 | 0 | 0 | 0 | 1 | 0 | 0 |
| "Taxon I" | 0 | 0 | 0 | 0 | 0 | 0 | 0 | 0 | 0 | 0 | 0 | 0 | 0 | 0 | 0 | 0 | 0 | 2 | 0 |

**Table S3:**
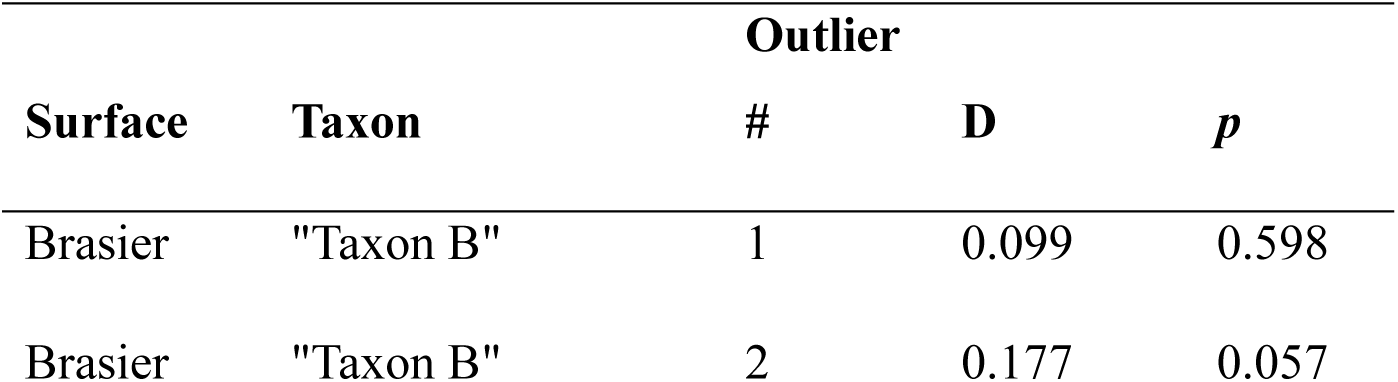
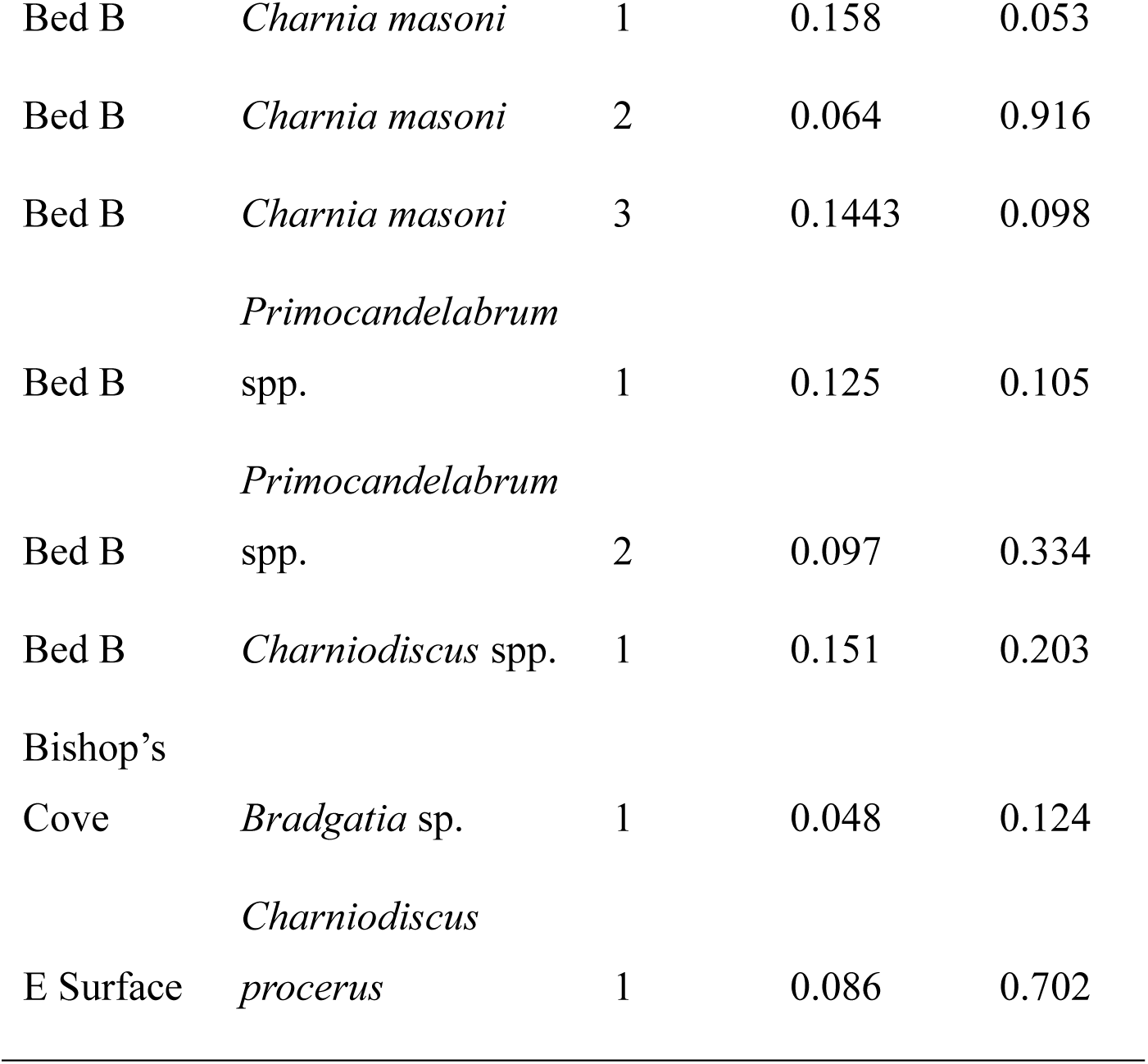
Kolmogorov-Smirnov test results for outsized specimens on Brasier and Bed B surfaces. Note the H26 outsized *Primocandelabrum* sp. specimen was not included here because the sample size of that population was too small.

## References

Aragonés Suarez, P., & Leys, S. P. (2022). The sponge pump as a morphological character in the fossil record. Paleobiology, 48(3), 446–461. 10.1017/pab.2021.43

Bače, R., Svoboda, M., Janda, P., Morrissey, R. C., Wild, J., Clear, J. L., Čada, V., & Donato, D. C. (2015). Legacy of Pre-Disturbance Spatial Pattern Determines Early Structural Diversity following Severe Disturbance in Montane Spruce Forests. PLOS ONE, 10(9), e0139214. 10.1371/journal.pone.0139214

Baddeley, A. (2015). Spatial point patterns: Methodology and applications with R (1st ed.). CRC Press.

Baddeley, A. (2024). *spatstat: Spatial Point Pattern Analysis, Model-Fitting, Simulation, Tests* (Version 3.1-1) [R]. CRAN. https://CRAN.R-project.org/package=spatstat

Bamforth, E. L., & Narbonne, G. M. (2009). New ediacaran rangeomorphs from Mistaken Point, Newfoundland, Canada. Journal of Paleontology, 83(6), 897–913. 10.1666/09-047.1

Bamforth, E. L., Narbonne, G. M., & Anderson, M. M. (2008). Growth and Ecology of a Multi-branched Ediacaran Rangeomorph from the Mistaken Point Assemblage, Newfoundland. Journal of Paleontology, 82(4), 763–777. 10.1666/07-112.1

Barnes, D. K. A., & Conlan, K. E. (2007). Disturbance, colonization and development of Antarctic benthic communities. Philosophical Transactions of the Royal Society B: Biological Sciences, 362(1477), 11–38. 10.1098/rstb.2006.1951

Benus, A. P. (1988). Faunas and depositional environments of the Upper Precambrian through lower Cambrian southeastern Newfoundland. 463, 18–52.

Bigham, K. T., Rowden, A. A., Leduc, D., & Bowden, D. A. (2021). Review and syntheses: Impacts of turbidity flows on deep-sea benthic communities. Biogeosciences, 18(5), 1893–1908. 10.5194/bg-18-1893-2021

Boddy, C. E., Mitchell, E. G., Merdith, A., & Liu, A. G. (2022). Palaeolatitudinal distribution of the Ediacaran macrobiota. Journal of the Geological Society, 179(1), jgs2021–030. 10.1144/jgs2021-030

Brasier, M. D., Antcliffe, J. B., & Liu, A. G. (2012). The architecture of Ediacaran Fronds. Palaeontology, 55(5), 1105–1124. 10.1111/j.1475-4983.2012.01164.x

Brunner, O., Chen, C., Giguère, T., Kawagucci, S., Tunnicliffe, V., Watanabe, H. K., & Mitarai, S. (2022). Species assemblage networks identify regional connectivity pathways among hydrothermal vents in the Northwest Pacific. Ecology and Evolution, 12(12), e9612. 10.1002/ece3.9612

Burt, A. J., Vogt-Vincent, N., Johnson, H., Sendell-Price, A., Kelly, S., Clegg, S. M., Head, C., Bunbury, N., Fleischer-Dogley, F., Jeremie, M.-M., Khan, N., Baxter, R., Gendron, G., Mason-Parker, C., Walton, R., & Turnbull, L. A. (2024). Integration of population genetics with oceanographic models reveals strong connectivity among coral reefs across Seychelles. Scientific Reports, 14(1), 4936. 10.1038/s41598-024-55459-x

Clapham, M. E., Narbonne, G. M., & Gehling, J. G. (2003). Paleoecology of the oldest known animal communities: Ediacaran assemblages at Mistaken Point, Newfoundland. Paleobiology, 29(4), 527–544. 10.1666/0094-8373(2003)029<0527:POTOKA>2.0.CO;2

Darroch, S. A. F., Laflamme, M., & Clapham, M. E. (2013). Population structure of the oldest known macroscopic communities from Mistaken Point, Newfoundland. Paleobiology, 39(4), 591–608. 10.1666/12051

Delahooke, K. M., Liu, A. G., Stephenson, N. P., & Mitchell, E. G. (2024). ‘Conga lines’ of Ediacaran fronds: Insights into the reproductive biology of early metazoans. Royal Society Open Science, 11(5), 231601. 10.1098/rsos.231601

Diggle, P. J. (2003). Statistical Analysis of Spatial and Spatio-Temporal Point Patterns (1st ed.). CRC Press.

Diggle, P. J., & Gratton, R. J. (1984). Monte Carlo Methods of Inference for Implicit Statistical Models. Journal of the Royal Statistical Society Series B; Statistical Methodology, 46, 193–212. http://www.jstor.org/stable/2345504

Dunn, F. S., Kenchington, C. G., Parry, L. A., Clark, J. W., Kendall, R. S., & Wilby, P. R. (2022). A crown-group cnidarian from the Ediacaran of Charnwood Forest, UK. Nature Ecology & Evolution, 6(8), 1095–1104. 10.1038/s41559-022-01807-x

Dunn, F. S., Liu, A. G., Grazhdankin, D. V., Vixseboxse, P., Flannery-Sutherland, J., Green, E., Harris, S., Wilby, P. R., & Donoghue, P. C. J. (2021). The developmental biology of *Charnia* and the eumetazoan affinity of the Ediacaran rangeomorphs. Science Advances, 7(30), eabe0291. 10.1126/sciadv.abe0291

Eden, R., Manica, A., & Mitchell, E. G. (2022). Metacommunity analyses show an increase in ecological specialisation throughout the Ediacaran period. PLOS Biology, 20(5), e3001289. 10.1371/journal.pbio.3001289

Gaylord, B., Reed, D. C., Raimondi, P. T., Washburn, L., & McLean, S. R. (2002). A physically based model of macroalgal spore dispersal in the wave and current-dominated nearshore. Ecology, 83(5), 1239–1251. 10.1890/0012-9658(2002)083[1239:APBMOM]2.0.CO;2

Getzin, S., Dean, C., He, F., A. Trofymow, J., Wiegand, K., & Wiegand, T. (2006). Spatial patterns and competition of tree species in a Douglas-fir chronosequence on Vancouver Island. Ecography, 29(5), 671–682. 10.1111/j.2006.0906-7590.04675.x

Getzin, S., Wiegand, T., Wiegand, K., & He, F. (2008). Heterogeneity influences spatial patterns and demographics in forest stands. Journal of Ecology, 96(4), 807–820. 10.1111/j.1365-2745.2008.01377.x

Grassle, J. F. (1977). Slow recolonisation of deep-sea sediment. Nature, 5595, 618–619.

Grazhdankin, D. V., Balthasar, U., Nagovitsin, K. E., & Kochnev, B. B. (2008). Carbonate-hosted Avalon-type fossils in arctic Siberia. Geology, 36(10), 803. 10.1130/G24946A.1

Griffiths, H. J., Whittle, R. J., & Mitchell, E. G. (2023). Animal survival strategies in Neoproterozoic ice worlds. Global Change Biology, 29(1), 10–20. 10.1111/gcb.16393

Hofmann, H. J., O’Brien, S. J., & King, A. F. (2008). Ediacaran biota on Bonavista Peninsula, Newfoundland, Canada. Journal of Paleontology, 82(1), 1–36. 10.1666/06-087.1

Ichaso, A. A., Dalrymple, R. W., & Narbonne, G. M. (2007). Paleoenvironmental and basin analysis of the late Neoproterozoic (Ediacaran) upper Conception and St. John’s groups, west Conception Bay, Newfoundland. Canadian Journal of Earth Sciences, 44(1), 25– 41. 10.1139/e06-098

Illian, J., Penttinen, A., Stoyan, H., & Stoyan, D. (2008). Statistical Analysis and Modelling of Spatial Point Patterns (1st ed.). Wiley.

Jacquemyn, H., Endels, P., Honnay, O., & Wiegand, T. (2010). Evaluating management interventions in small populations of a perennial herb *Primula vulgaris* using spatio-temporal analyses of point patterns. Journal of Applied Ecology, 47(2), 431–440. 10.1111/j.1365-2664.2010.01778.x

Jõgiste, K., Korjus, H., Stanturf, J. A., Frelich, L. E., Baders, E., Donis, J., Jansons, A., Kangur, A., Köster, K., Laarmann, D., Maaten, T., Marozas, V., Metslaid, M., Nigul, K., Polyachenko, O., Randveer, T., & Vodde, F. (2017). Hemiboreal forest: Natural disturbances and the importance of ecosystem legacies to management. Ecosphere, 8(2), e01706. 10.1002/ecs2.1706

Kenchington, C. G., Dunn, F. S., & Wilby, P. R. (2018). Modularity and Overcompensatory Growth in Ediacaran Rangeomorphs Demonstrate Early Adaptations for Coping with Environmental Pressures. Current Biology, 28(20), 3330–3336.e2. 10.1016/j.cub.2018.08.036

Khan, T. M., Griffiths, H. J., Whittle, R. J., Stephenson, N. P., Delahooke, K. M., Purser, A., Manica, A., & Mitchell, E. G. (2024). Network analyses on photographic surveys reveal that invertebrate predators do not structure epibenthos in the deep (∼2000m) rocky Powell Basin, Weddell Sea, Antarctica. Frontiers in Marine Science, 11, 1408828. 10.3389/fmars.2024.1408828

Kim, S., Wild, C., & Tilstra, A. (2022). Effective asexual reproduction of a widespread soft coral: Comparative assessment of four different fragmentation methods. PeerJ, 10, e12589. 10.7717/peerj.12589

Kolmogorov, A. (1933). Sulla determinazione empirica di una legge di distrubuzione. Giornale Dell’Istituto Italiano Degli Attuari, 4, 83–91.

Le Bas, M. J. (1984). Geological evidence from Leicestershire on the crust of southern Britain. Transactions of the Leicester Literary and Philosophical Society, 76, 54–67.

Liu, A. G., & Dunn, F. S. (2020). Filamentous Connections between Ediacaran Fronds. Current Biology, 30(7), 1322–1328.e3. 10.1016/j.cub.2020.01.052

Liu, A. G., Kenchington, C. G., & Mitchell, E. G. (2015). Remarkable insights into the paleoecology of the Avalonian Ediacaran macrobiota. Gondwana Research, 27(4), 1355–1380. 10.1016/j.gr.2014.11.002

Maldonado, M., & Uriz, M. J. (1999). Sexual propagation by sponge fragments. Nature, 398(6727), 476–476. 10.1038/19007

Mann, H. B., & Whitney, D. R. (1947). On a Test of Whether one of Two Random Variables is Stochastically Larger than the Other. The Annals of Mathematical Statistics, 18(1), 50– 60. 10.1214/aoms/1177730491

Matthews, J. J., Liu, A. G., Yang, C., McIlroy, D., Levell, B., & Condon, D. J. (2021). A Chronostratigraphic Framework for the Rise of the Ediacaran Macrobiota: New Constraints from Mistaken Point Ecological Reserve, Newfoundland. Geological Society of America Bulletin, 133(3).

McFadden, I. R., Bartlett, M. K., Wiegand, T., Turner, B. L., Sack, L., Valencia, R., & Kraft, N. J. B. (2019). Disentangling the functional trait correlates of spatial aggregation in tropical forest trees. Ecology, 100(3), e02591. 10.1002/ecy.2591

Millard, S. P. (2013). EnvStats: An R Package for Environmental Statistics. Springer. https://www.springer.com

Mitchell, E. G., Bobkov, N., Bykova, N., Dhungana, A., Kolesnikov, A. V., Hogarth, I. R. P., Liu, A. G., Mustill, T. M. R., Sozonov, N., Rogov, V. I., Xiao, S., & Grazhdankin, D. V. (2020). The influence of environmental setting on the community ecology of Ediacaran organisms. Interface Focus, 10(20190109). 10.1098/rsfs.2019.0109

Mitchell, E. G., & Harris, S. (2020). Mortality, Population and Community Dynamics of the Glass Sponge Dominated Community “The Forest of the Weird” From the Ridge Seamount, Johnston Atoll, Pacific Ocean. Frontiers in Marine Science, 7, 565171. 10.3389/fmars.2020.565171

Mitchell, E. G., Harris, S., Kenchington, C. G., Vixseboxse, P., Roberts, L., Clark, C., Dennis, A., Liu, A. G., & Wilby, P. R. (2019). The importance of neutral over niche processes in structuring Ediacaran early animal communities. Ecology Letters, 22(12), 2028– 2038. 10.1111/ele.13383

Mitchell, E. G., & Kenchington, C. G. (2018). The utility of height for the Ediacaran organisms of Mistaken Point. Nature Ecology & Evolution, 2(8), 1218–1222. 10.1038/s41559-018-0591-6

Mitchell, E. G., Kenchington, C. G., Harris, S., & Wilby, P. R. (2018). Revealing rangeomorph species characters using spatial analyses. Canadian Journal of Earth Sciences, 55(11), 1262–1270. 10.1139/cjes-2018-0034

Mitchell, E. G., Kenchington, C. G., Liu, A. G., Matthews, J. J., & Butterfield, N. J. (2015). Reconstructing the reproductive mode of an Ediacaran macro-organism. Nature, 524(7565), 343–346. 10.1038/nature14646

Mitchell, E. G., & Pates, S. (2025). From organisms to biodiversity: The ecology of the Ediacaran/Cambrian transition. Paleobiology, 1–24. 10.1017/pab.2024.21

Myrow, P. (1995). Neoproterozoic rocks of the Newfoundland Avalon Zone. Precambrian Research, 73(1–4), 123–136. 10.1016/0301-9268(94)00074-2

Narbonne, G. M. (2004). Modular Construction of Early Ediacaran Complex Life Forms. Science, 305(5687), 1141–1144. 10.1126/science.1099727

Narbonne, G. M. (2005). The Ediacara biota: Neoproterozoic Origin of Animals and Their Ecosystems. Annual Review of Earth and Planetary Sciences, 33(1), 421–442. 10.1146/annurev.earth.33.092203.122519

Noble, S. R., Condon, D. J., Carney, J. N., Wilby, P. R., Pharaoh, T. C., & Ford, T. D. (2015). U-Pb geochronology and global context of the Charnian Supergroup, UK: Constraints on the age of key Ediacaran fossil assemblages. Geological Society of America Bulletin, 127(1).

Nyström, M., & Folke, C. (2001). Spatial Resilience of Coral Reefs. Ecosystems, 4(5), 406– 417. 10.1007/s10021-001-0019-y

O’Brien, S. J., & King, A. F. (2005). Late neoproterozoic (Ediacaran) stratigraphy of Avalon zone sedimentary rocks, Bonavista Peninsula, Newfoundland. Current Research.

O’Brien, S. J., Wardle, R. J., & King, A. F. (1983). The Avalon Zone: A Pan-African terrane in the Appalachian Orogen of Canada. Geological Journal, 18(3), 195–222. 10.1002/gj.3350180302

O’Connell, B., McMahon, W. J., Nduutepo, A., Pokolo, P., Mocke, H., McMahon, S., Boddy, C. E., & Liu, A. G. (2024). Transport of ‘Nama’-type biota in sediment gravity and combined flows: Implications for terminal Ediacaran palaeoecology. Sedimentology, sed.13239. 10.1111/sed.13239

Okubo, N., Motokawa, T., & Omori, M. (2007). When fragmented coral spawn? Effect of size and timing on survivorship and fecundity of fragmentation in *Acropora formosa*. Marine Biology, 151(1), 353–363. 10.1007/s00227-006-0490-2

Pasinetti, G., & McIlroy, D. (2023). Palaeobiology and taphonomy of the rangeomorph *Culmofrons plumosa*. Palaeontology, 66(4), e12671. 10.1111/pala.12671

Pérez-Hernández, J., & Gavilán, R. G. (2021). Impacts of Land-Use Changes on Vegetation and Ecosystem Functioning: Old-Field Secondary Succession. Plants, 10(5), 990. 10.3390/plants10050990

Pharaoh, T. C., Webb, P. C., Thorpe, R. S., & Beckinsale, R. D. (1987). Geochemical Evidence for the Tectonic Setting of Late Proterozoic Volcanic Suites in Central England. Geological Society, London, Special Publications, 33(1), 541–552. 10.1144/GSL.SP.1987.033.01.36

Potthoff, M., Johst, K., & Gutt, J. (2006). How to survive as a pioneer species in the Antarctic benthos: Minimum dispersal distance as a function of lifetime and disturbance. Polar Biology, 29(7), 543–551. 10.1007/s00300-005-0086-1

Presley, S. J., Higgins, C. L., & Willig, M. R. (2010). A comprehensive framework for the evaluation of metacommunity structure. Oikos, 119(6), 908–917. 10.1111/j.1600-0706.2010.18544.x

R Core Team. (2023). R: A Language and Environment for Statistical Computing (Version 4.2.2) [Computer software].

Raventós, J., Wiegand, T., & Luis, M. D. (2010). Evidence for the spatial segregation hypothesis: A test with nine-year survivorship data in a Mediterranean shrubland. Ecology, 91(7), 2110–2120. 10.1890/09-0385.1

Runnegar, B. (2022). Following the logic behind biological interpretations of the Ediacaran biotas. Geological Magazine, 159(7), 1093–1117. 10.1017/S0016756821000443

Shen, G., Yu, M., Hu, X.-S., Mi, X., Ren, H., Sun, I.-F., & Ma, K. (2009). Species–area relationships explained by the joint effects of dispersal limitation and habitat heterogeneity. Ecology, 90(11), 3033–3041. 10.1890/08-1646.1

Smirnov, N. (1948). Table for estimating the goodness of fit of empirical distributions. Annals of Mathematical Statistics, 19, 279–281.

Smith, K. E., Aronson, R. B., Steffel, B. V., Amsler, M. O., Thatje, S., Singh, H., Anderson, J., Brothers, C. J., Brown, A., Ellis, D. S., Havenhand, J. N., James, W. R., Moksnes, P. - O., Randolph, A. W., Sayre-McCord, T., & McClintock, J. B. (2017). Climate change and the threat of novel marine predators in Antarctica. Ecosphere, 8(11), e02017. 10.1002/ecs2.2017

Sperling, E. A., Peterson, K. J., & Laflamme, M. (2011). Rangeomorphs, *Thectardis* (Porifera?) and dissolved organic carbon in the Ediacaran oceans: Rangeomorphs, Thectardis and DOC. Geobiology, 9(1), 24–33. 10.1111/j.1472-4669.2010.00259.x

Stephenson, N. P., Delahooke, K. M., Barnes, N., Rideout, B. W. T., Kenchington, C. G., Manica, A., & Mitchell, E. G. (2024). Morphology shapes community dynamics in early animal ecosystems. Nature Ecology & Evolution, 8(7), 1238–1247. 10.1038/s41559-024-02422-8

Thatje, S. (2012). Effects of Capability for Dispersal on the Evolution of Diversity in Antarctic Benthos. Integrative and Comparative Biology, 52(4), 470–482. 10.1093/icb/ics105

Thomas, M. (1949). A Generalization of Poisson’s Binomial Limit For use in Ecology. Biometrika, 36(1), 18–25.

Thrush, S. F., Hewitt, J. E., Cummings, V. J., Norkko, A., & Chiantore, M. (2010). β-Diversity and Species Accumulation in Antarctic Coastal Benthos: Influence of Habitat, Distance and Productivity on Ecological Connectivity. PLoS ONE, 5(7), e11899. 10.1371/journal.pone.0011899

Velázquez, E., Martínez, I., Getzin, S., Moloney, K. A., & Wiegand, T. (2016). An evaluation of the state of spatial point pattern analysis in ecology. Ecography, 39(11), 1042–1055. 10.1111/ecog.01579

Wiegand, T. (2014). Programita (Version February 2014) [Computer software].

Wiegand, T. (2018). Programita (Version November 2018) [Computer software].

Wiegand, T., & Moloney, K. A. (2014). Handbook of Spatial Point-Pattern Analysis in Ecology (1st ed.). CRC Press.

Wilby, P. R., Kenchington, C. G., & Wilby, R. L. (2015). Role of low intensity environmental disturbance in structuring the earliest (Ediacaran) macrobenthic tiered communities. Palaeogeography, Palaeoclimatology, Palaeoecology, 434, 14–27. 10.1016/j.palaeo.2015.03.033

Wild, J., Kopecký, M., Svoboda, M., Zenáhlíková, J., Edwards-Jonášová, M., & Herben, T. (2014). Spatial patterns with memory: Tree regeneration after stand-replacing disturbance in *Picea abies* mountain forests. Journal of Vegetation Science, 25(6), 1327–1340. 10.1111/jvs.12189

Wood, D. A., Dalrymple, R. W., Narbonne, G. M., Gehling, J. G., & Clapham, M. E. (2003). Paleoenvironmental analysis of the late Neoproterozoic Mistaken Point and Trepassey formations, southeastern Newfoundland. Canadian Journal of Earth Sciences, 40(10), 1375–1391. 10.1139/e03-048

Wu, C., Pang, K., Chen, Z., Wang, X., Zhou, C., Wan, B., Yuan, X., & Xiao, S. (2024). The rangeomorph fossil *Charnia* from the Ediacaran Shibantan biota in the Yangtze Gorges area, South China. Journal of Paleontology, 98(2), 232–248. 10.1017/jpa.2022.97

